# Socio-ecological context modulates significance of territorial contest competition in *Drosophila prolongata*

**DOI:** 10.1101/2024.04.26.587716

**Authors:** Alessio N. De Nardo, Broti Biswas, Jhoniel Perdigón Ferreira, Abhishek Meena, Stefan Lüpold

## Abstract

The intensity and direction of sexual selection is intricately linked to the social and ecological context. Both operational sex ratios (OSRs) and population densities can affect the ability of males to monopolize resources and mates, and thus the form and intensity of sexual selection on them. Here, we studied how the mating system of the promiscuous and strongly sexually dimorphic fruit fly *Drosophila prolongata* responds to changes in the OSR and population density. We recorded groups of flies over five days and quantified territory occupancy, mating success, and competitive fertilization success. Although sexual selection was stronger under male-biased than even OSRs but unrelated to density, realized selection on morphological traits was higher under even OSRs and increased with density. Larger and more territorial males achieved both higher mating success and competitive fertilization success, but only under even OSRs. Our combined results also support a shift in the mating system from territorial contest competition to scramble competition under male-biased OSRs and potentially at low density, where there was no clear contribution of the measured traits to reproductive success. Our study emphasizes the limitations of traditional selection metrics and the role of the socio-ecological context in predicting adaptation to a changing environment.

**Significance:** Mating systems are complex and dynamic, adapting to ongoing ecological change. Studies often assume that changes in the socio-ecological context alter the intensity of sexual selection on traits indicating individual fitness, but our work on *Drosophila prolongata* challenges this view. By manipulating operational sex ratio and population density, and jointly investigating territorial behavior and both pre- and post-mating reproductive success over several days, we reveal the plastic mating strategies in this fly. This dynamism underscores the limitations of static classifications and the importance of studying selection across diverse socio-ecological contexts. This broader perspective advances our understanding of the tight connections between environmental change, population demographics, and the evolutionary process.

## Introduction

Sexual selection, driven by competition for mates and/or fertilization (1), shapes heritable traits and behaviors that are directly related to reproductive success (2). While sexual selection can operate in both sexes, it tends to be stronger on males than females (2, 3). Much of this asymmetry in sexual selection is attributed to the typically greater investments per offspring by females compared to males, which reduces their reproductive potential and intensifies the competition among males over the limited reproductive opportunities (4, 5). The direction and intensity of sexual selection between the sexes can be influenced by the ecological context. For example, food scarcity in katydid bush crickets shifts the typical sex roles to females competing for access to the nutritious spermatophores provided by males (6).

One important ecological parameter affecting the intensity of sexual selection is the distribution and defensibility of resources, which can shape the local population composition and density. For example, variation in the operational sex ratio (OSR; i.e., the ratio of sexually receptive males to sexually receptive females (7)) can affect the availability of potential mates. A skewed OSR can lead to monopolization of mating opportunities by a few successful individuals of the competing sex (7). Male-biased OSRs in particular can thus drive selection on traits that are advantageous in competition, including larger body size, the growth of weaponry, or elaborate ornamentation favored by females (7–9).

A meta-analysis across 62 empirical studies of 58 different species generally supports theoretical predictions that OSRs can drive the intensity of sexual selection (10). However, studies exploring selection on specific phenotypic traits often find no difference in the strength of selection between different OSRs (e.g., on body size (11–13)), or they show increased selection intensity in even or female-biased populations (14–16). For example, in the common lizard (*Lacerta vivipara*), females favor small males under male-biased OSRs as large males leave more bite scars on them during mating (14, 17). Overall, the effect of the OSR on the intensity of sexual selection appears to vary considerably across species and mating systems (18, 19).

Apart from mate competition, OSRs can also influence post-mating processes, such as competitive fertilization (20–22; but see 23), ultimately impacting aspects like ejaculate transfer or mating duration (24). Males tend to invest more in ejaculate production when exposed to a higher risk or intensity of sperm competition (20), often induced by changes in the perceived OSR (25–27). Thus, traits favored in the post-mating phase, such as enhanced ejaculate allocation (28) or larger testes (29) are expected to be under stronger selection when OSRs are male-biased, yet studies on the latter are inconclusive (30–36). In a literature review (30), only one of four studies found a significant effect of OSR on testes size. This one study showed an increase in testes size for male *Drosophila melanogaster* evolving under an extreme female bias (10:1), suggesting selection against sperm limitation rather than sperm competition (31).

In addition to the OSR, competition for mates can also vary with population density. Theoretical work suggests that higher population densities increase the encounter rate both between potential mates and competitors, which can in turn intensify sexual selection (37). By contrast, the scope for sexual selection may be more limited at low densities, where infrequent mate encounters may lower a female’s mate discrimination due to higher sampling costs (37–39), and rare encounters among males also limit the potential for mate competition (37). Empirical studies on these patterns, however, have yielded mixed results (9, 40–48). Whereas some studies point toward more intense sexual selection at higher densities (40–42), others report the opposite (44–46) or no relationship (9, 47, 48). One possible reason for these contrasting results is that the traits examined in these studies are under pre-mating sexual selection, which might trade off with traits under post-mating sexual selection in a density-dependent manner (36, 49). For example, higher population densities might increase the relative importance of post-mating competition if increasing costs of monopolization lower the effectiveness of aggressive strategies (36, 37, 45, 50, 51). Such a trend is suggested across frogs and toads, where the trade-off between forelimb muscles and testes size is biased more strongly toward the latter as population densities increase and competitive fertilization becomes more difficult for males to prevent (36).

Sexual selection and male reproductive tactics may also be influenced by the distribution of resources (7). While evenly distributed resources limit the opportunity for monopolization, highly competitive males may gain control over spatially concentrated resources (7). These males may then become more attractive to potential mates for their ability to defend a resource or for the benefit the resource provides, tying mating success to resource monopolization (52–54). For example, in antlered flies, *Phytalmia spp.,* males fight over oviposition sites to which females then gain access through copulation (55). Therefore, females indirectly choose mates with large antlers, which in turn benefit males in these contests (56). However, reproductive tactics may vary between, or temporally within, individuals (57), mediated by the number of individuals competing for a resource and how well it can be defended (58, 59). Males may follow a territorial or a non-territorial strategy, depending on the relative benefits of each for a given competitor density (45, 51, 60) or their own resource-holding potential (52), or they may employ a flexible strategy (61, 62). Similarly, if the population density or the physical condition of a male limit his reproductive success by courting or competing for females, he may adopt an alternative tactic, such as attempting to steal copulations from courting males (63–66).

Although OSRs and population densities have been shown to shape sexual selection and mating systems, both separately and jointly (e.g., 8, 9, 40, 41, 44, 49, 67), it remains unknown how these factors combined drive both pre- and post-mating sexual selection, particularly in the context of uneven resource distribution. Here, we investigated these factors in a comprehensive laboratory experiment using the fruit fly *Drosophila prolongata.* This South-Asian species exhibits conspicuous male-biased sexual dimorphism both in body size and in foreleg size and coloration (68), while also showing resource-defense behavior when high-quality resources are concentrated (69). Males use their forelegs in both fights against competitors (70, 71) and elaborate courtship behaviors (68). These behaviors also include “leg vibration”, a species-specific behavior that involves a male stimulating the female abdomen and is often displayed immediately before copulation (71), likely to overcome female remating resistance (72, 73). Leg vibration, however, can also trigger an alternative reproductive tactic by a nearby male, namely interception of the female (66). With highly polygamous females (72), this species is ideal for studies of context-dependent pre- and post-mating sexual selection.

Here, we monitored over a five-day interaction period each of 30 groups of *D. prolongata* that varied in their sex ratio (1:1 or 3:1 males:females) and density (12, 16, or 20 flies). Over five days, we quantified the distribution of flies around a centrally located resource, their pre-mating behaviors, individual copulations, and molecular parentage, which provided a uniquely comprehensive understanding of reproductive dynamics in this species. We predicted the intensity of sexual selection on males to increase (i) under a male-biased OSR and (ii) with population density, translating into intensifying selection on traits mediating resource defense and mate competition, such as male body size or foreleg length. We also predicted that (iii) male-biased OSRs and high densities should increase the relative importance of post-relative to pre-mating sexual selection.

## Results and Discussion

### Descriptive statistics and skew in reproductive success

Across all 180 females and 300 males, we observed 754 matings, with only three females (1.6%) and 83 males (27%) never mating. Most copulations occurred on the first experimental day (67.6%), with another 13%, 10.5%, 5.7%, and 3.2% of matings on days 2−5, respectively. Across all five days, females mated 0−13 times (mean±SD = 3.95±2.52), with up to 7 different sires (2.36±1.39; Fig. 1). Males gained 0−14 matings (2.50±2.6) across up to 9 females (1.47±1.59; Fig. 1). Varying numbers of possible matings between treatments hampered direct comparisons of absolute mating numbers. However, relative male mating success was more heavily skewed under male-biased compared to even OSRs despite a similar mean success rate (Fig. S1), suggesting more intense sexual selection under a male-biased OSR. These differences were more subtle between densities, but generally suggested weaker sexual selection at low compared to medium or high densities (Fig. S2).

**Figure 1:**
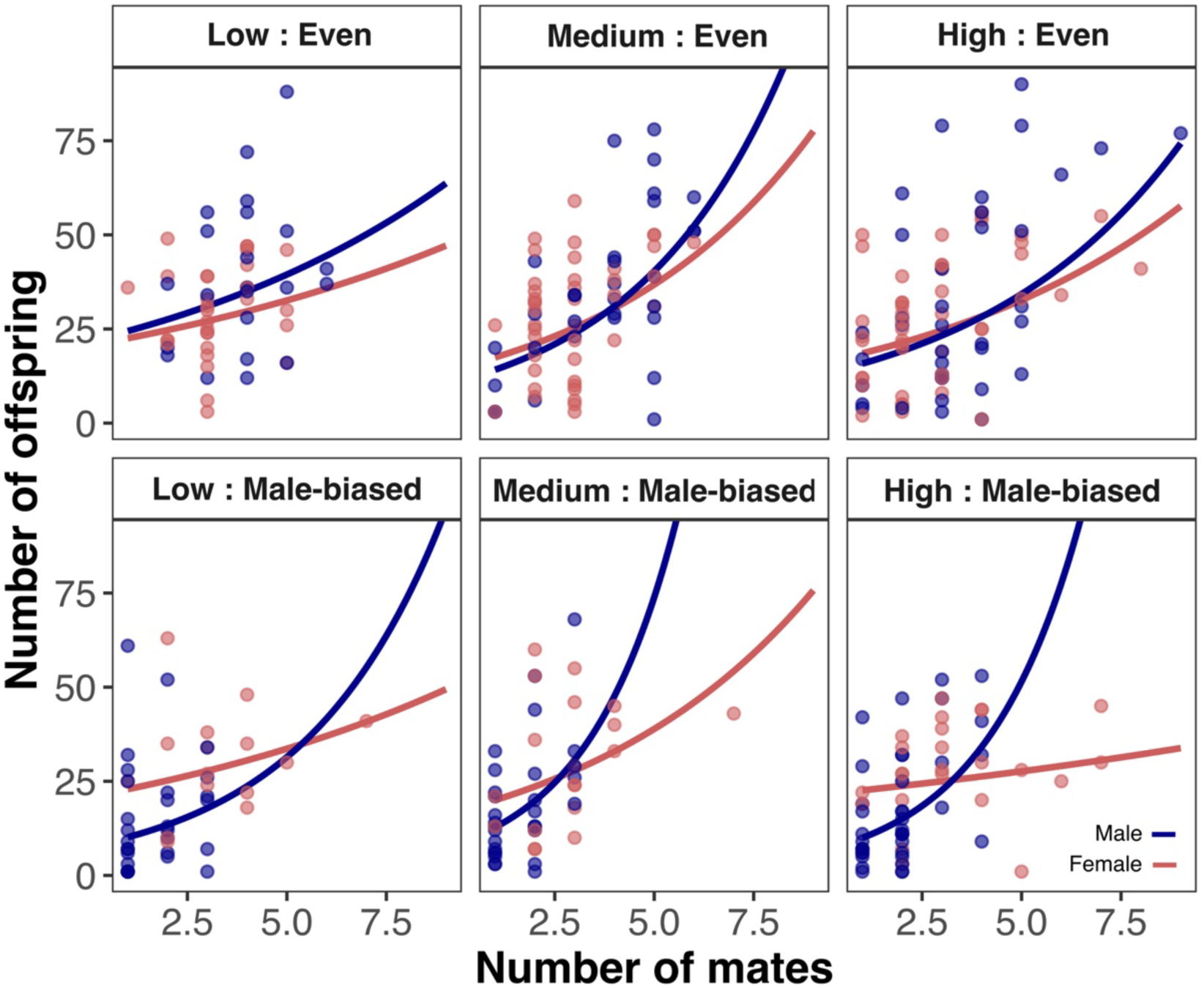
Bateman gradients for males (blue) and females (red) for each combination of density (low, medium, high) and OSR (even, male-biased) and density.

### Potential and realized sexual selection

To quantify the selection dynamics in *D. prolongata* under different population compositions, we measured Bateman gradients and the opportunity for sexual selection (*I*_s_) as two commonly used estimates for the intensity of sexual selection (10). Whereas the former represents the slope between individuals’ number of mates and resulting offspring (i.e., the fitness advantage gained by additional matings (74)), the latter is based on the sex-specific variances in reproductive success (i.e., the opportunity for selection to act on variation in reproductive success (75)). The Bateman gradients for both sexes were positive (Fig. 1; males: *X*^2^_1_ = 63.65, *n* = 193, *P* < 0.001, Table S1; females: *X*^2^_1_ = 24.1, *n* = 175, *P* < 0.001, Table S2), but more pronounced in males (*X*^2^_1_ = 7.42, *P* = 0.006, Table S3a). The disparity in Bateman gradients between the sexes was greater in the male-biased populations (*X*^2^_1_ = 5.6, *n =* 368, *P* = 0.018) but unaffected by group density (*X*^2^_2_ = 0.23, *P* = 0.89, Table S3b). Similarly, *I*_s_ was higher for males compared to females in all treatments, but particularly pronounced in the male-biased populations (6.2× higher for males) compared to the even OSR populations (2.7× higher for males). Population density again had no significant effect (Table S4, Fig. S3). These findings combined suggest that sexual selection is more pronounced on males than females, and its intensity for males is amplified under a male-biased OSR, as previously shown for a diversity of taxa (10).

When comparing selection on specific male traits, results were more complex. For example, we found positive selection gradients on thorax, foreleg and wing length, and composite body size (principal component 1, PC1; see *Methods*) under even OSR, but only on wing length in male-biased populations (Fig. 2, Table S5). Also, contrary to expectations based on *I*_s_, positive selection on all measured traits appeared to intensify with density (Fig. 2, Table S6). This increasing selection on secondary sexual traits with density aligns with theory (37), yet has been reported for only few species (42, 76, 77; but see refs. 45, 78). The changing pattern of selection on phenotypic traits across OSRs, however, contradicts both our predictions and insights gained from the above Bateman gradients and *I*_s_.

**Figure 2:**
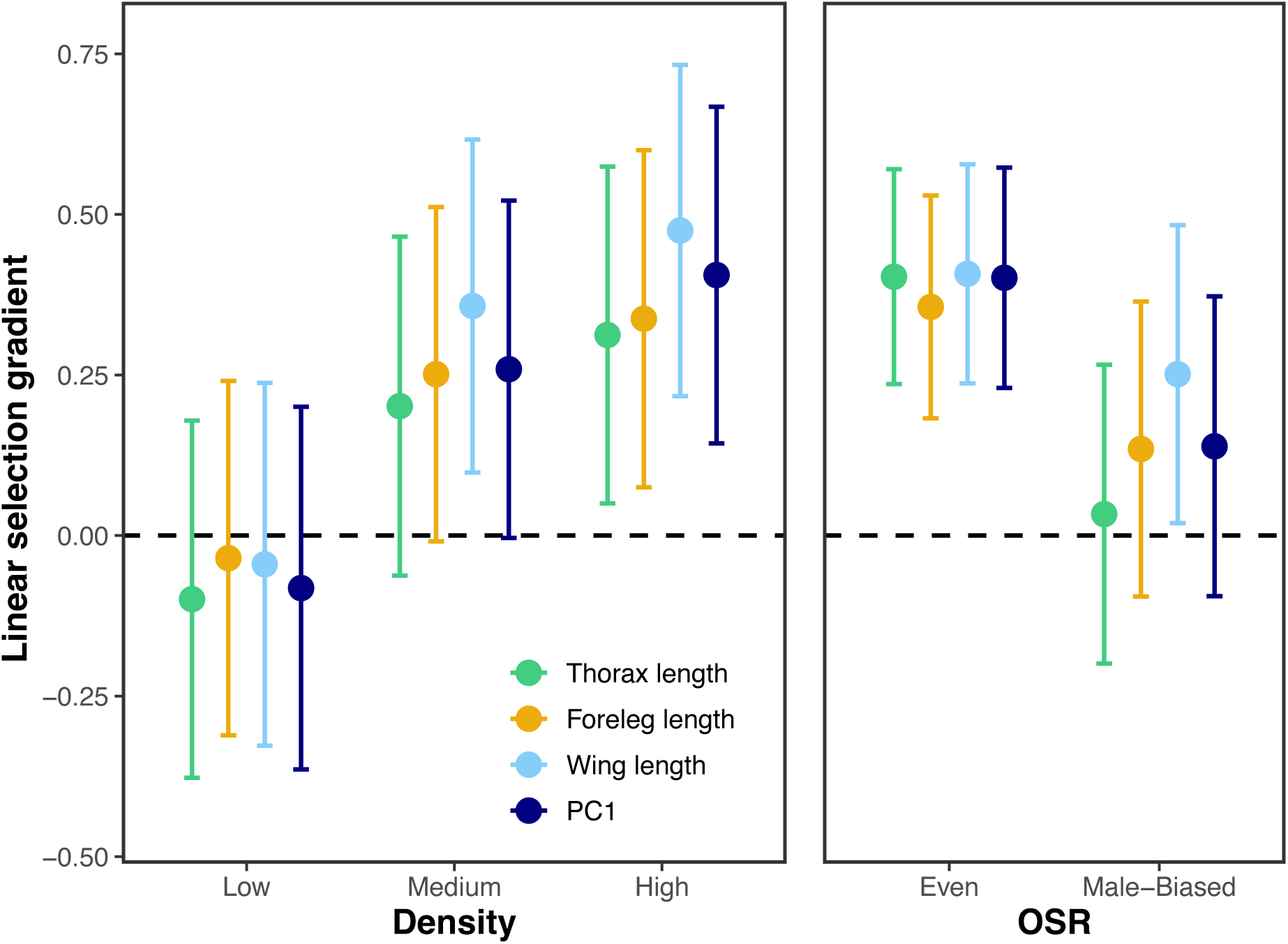
Linear selection gradients with 95% confidence intervals on male morphological traits under different levels of population density and OSRs.

The lack of positive selection on putative sexually selected traits in male-biased populations might simply indicate insufficient statistical power or that other, unmeasured behavioral or morphological traits, or traits under post-mating selection would be favored when mates are more difficult to monopolize (2, 79, 80). However, our findings might also reveal the limitations of using metrics like Bateman gradients, *I*_s_, or the OSR to predict realized selection on specific sexually selected traits, as previously emphasized (18). As our results confirm, these common measures of the intensity of sexual selection do not necessarily correlate with the observed selection on traits and merely set an upper limit to the potential of sexual selection (18). These results underscore the need to integrate different measures to gain a more complete understanding of the dynamics of a mating system. Despite their limitations, Bateman gradients or *I*_s_ remain valuable tools for numerically characterizing mating systems. They consider all types of sexual selection (e.g. pre- and post-mating, intra- and intersexual), acting on any relevant morphological or behavioral traits (81). Thus, they accurately capture variation in fitness and allow for comparisons between sexes and populations (81). For example, we showed that reproductive success is more skewed toward a few successful males in male-biased populations than under even OSR. The trait-specific selection gradients indicate, however, that traits selected in male-biased populations are not the morphological traits expected to be involved in physical contest such as resource defense (e.g., forelegs). These insights emphasize the complex interplay between OSR, density, and sexual selection in shaping mating system dynamics and trait evolution in this species.

### Territorial behavior and mating success

To investigate how the territorial behavior previously reported in this species (69) manifests in different group compositions, a resource-dense area (RDA) was established in the center of each experimental unit. This RDA, consisting of highly nutritious yeast paste, covered approximately 16% of the otherwise nutrient-poor arena. Regardless of population composition, males spent more time in the RDA than females (35.6% vs. 12.3%; *X*^2^_1_ = 671.6, *P* < 0.001, *n* = 493 (303 males, 190 females), Table S7), with greater variance in males (*σ*^2^ = 0.016) than in females (*σ*^2^ = 0.005). Hence, the RDA appeared to be highly attractive to males, but access was unequal among them as reflected by their varying time spent there (indicating “territorial success”). In addition, individuals of both sexes spent less time (on average) in the RDA at higher densities (*X*^2^_1_ = 22.0, *P* < 0.001) and under male-biased OSRs (*X*^2^_1_ = 21.0, *P* < 0.001), suggesting intensifying competition for this resource. Although females spent considerably less time in the RDA overall, 69.4% of all matings started there. These matings were also more likely to involve aggressive male-male behavior than matings outside this area (*z* = 2.54, *P* = 0.011, *n* = 716 matings across 212 males and 187 females). Both sexes spent more time in the RDA on the first day of recording, when most matings occurred, compared to the subsequent days (*X*^2^_4_ = 313.724, *P* < 0.001, *n =* 2379 across 5 days and 492 individuals; Fig. S4, Table S8). Interestingly, body size (PC1) had no effect on territorial occupancy on the first day, but did so on all following days (interaction: *X*^2^_1_ = 5.05, *n* = 1445, *P* = 0.02, Table S9), suggesting that males might initially need time to establish dominance hierarchies and resource access, with large males ultimately being more successful (69, 82). Additionally, given the females’ heightened receptivity on day one, following females around may have been advantageous to staying in the RDA, also for large males. Indeed, matings were, if anything, less likely to start in the RDA on the first (66.9%) compared to later days (73.8%; *z* = 1.78, *P* = 0.075).

Amino & Matsuo (69) report a hierarchy in pairs of males, with one male clearly dominating territory occupancy, particularly when the area is small and easily defendable. Although their findings do not directly demonstrate mating success by territory occupancy, a strong correlation between territory occupancy and courtship rate (69), as well as a mating advantage for winners of contests over territories (83), have been reported for *D. prolongata*. In our study, the RDA was comparable to the larger of the two areas used by Amino & Matsuo (69), but with more males and so reduced ability to monopolize the contested area. Nevertheless, territorial control influenced mating success, albeit with greater complexity at higher numbers of competitors: Compared to 72.5% at an even OSR, only 64.6% (*z* = 1.85, *P* = 0.065) of matings were initiated in the RDA at a male-biased OSR. Interestingly, although most matings were initiated in the RDA at both OSRs, overall territory occupancy determined mating success only in even (*t* = 2.4, *P* = 0.019) but not male-biased populations (*t* = 0.24, *P* = 0.8; OSR×territoriality: *t* = 1.7, *P* = 0.09, *n* = 290, controlling for PC1; Fig. 3, Table S10a-b). Similarly, PC1 was the main contributor to mating success under even (*t* = 3.7, *P* < 0.001) but not male-biased OSRs (*t* = 1.1, *P* = 0.26, OSR×PC1: *t* = 2.15, *P* = 0.03*; n =* 290, Fig. 3, Table S10a-b). In addition to these results, the occurrence of courtship behavior before mating depended on body size and the OSR (PC1×OSR: *t* = 2.93, *P* = 0.003, *n* = 760, Fig. 3, Table S14a), in that large males showed more courtship behavior in the even OSR (*z* = 3.20, *P* = 0.001) but not in the male-biased OSR (*z* = −1.27, *P* = 0.20). These results align with the selection gradients above, but additionally illustrate the changing dynamics of sexual selection in response to varying levels of competition.

**Figure 3:**
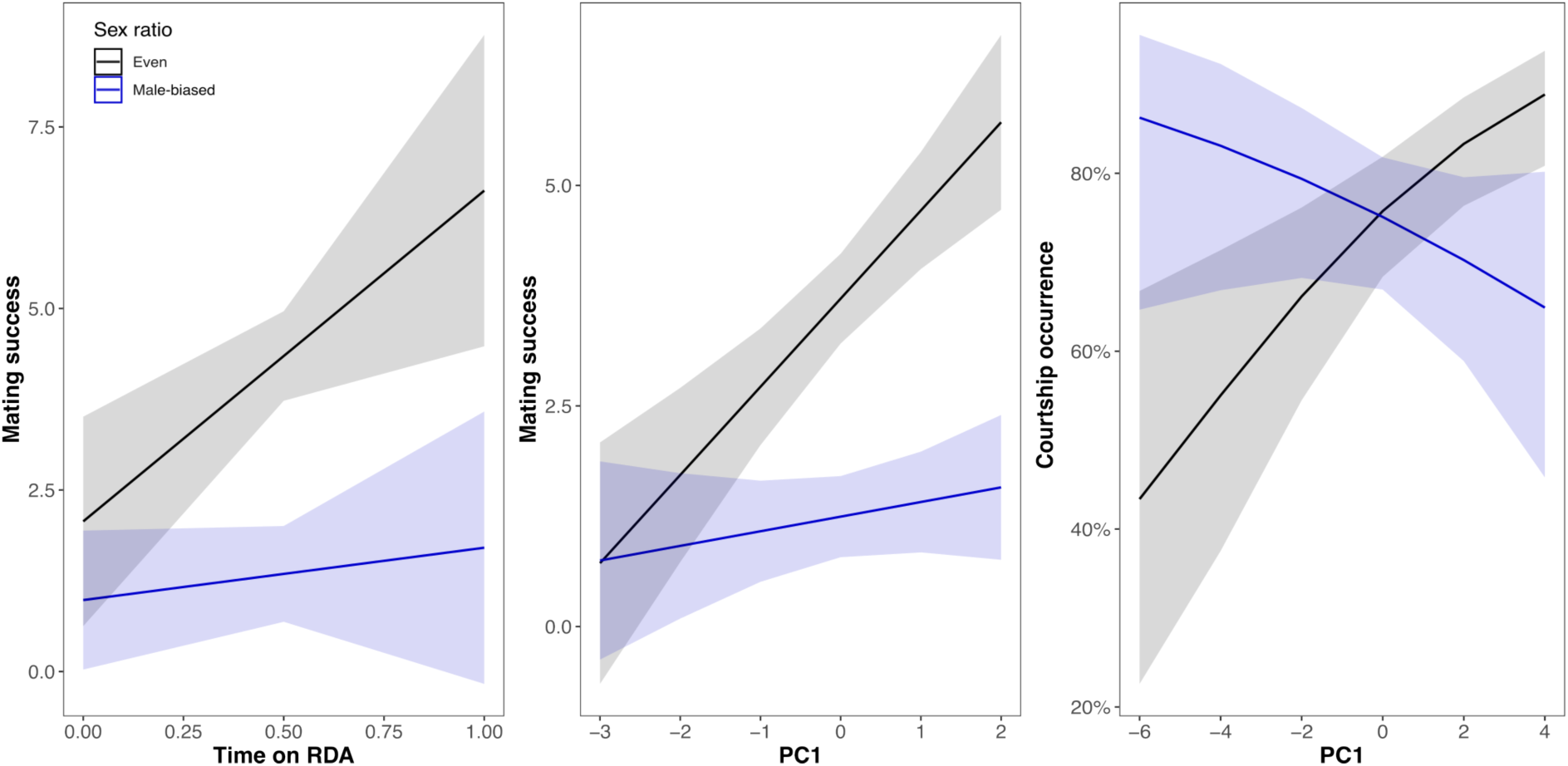
The effect of territoriality (time on the resource-dense area; left), and PC1 (middle) on mating success depends on the OSR. The effect of PC1 on courtship occurrence also depends on the OSR (right).

Our findings support the notion that when competition intensifies (e.g., strongly male-biased OSR), monopolization of resources becomes increasingly difficult, potentially shifting mating strategies (36, 37, 49, 50, 64). However, we neither observed an increase in sneak matings at higher densities nor under male-biased OSRs (both |*z*| < 1.2, *P* > 0.26, *n* = 716, Table S12). In fact, leg vibration, an auditory cue used by sneaky males to intercept a mating attempt, even tended to decrease at high densities (*z* = −1.7, *P* = 0.088) and in male-biased OSRs (*z* = 1.89, *P* = 0.059, *n* = 716, Table S11). Courtship was observed less frequently when matings occurred in the RDA (*z* = −3.3, *P* < 0.001, Table S14), possibly due to a high local density of competitors and so higher risk of becoming the target of aggression (*z* = 2.54, *P* = 0.011, *n* = 716, Table S13) or interception (*z* = 2.31, *P* = 0.021, *n* = 716, Table S13). Similarly, leg vibration was observed less often before matings in the RDA (*z* = −1.81, *P* = 0.07, *n* = 716, Table S11). Large males were more likely to gain matings following leg vibration (*z* = 2.89, *P* = 0.004, *n* = 716), whereas those acquiring matings by interception tended to be smaller males (*t* = −2.03, *P* = 0.042, *n* = 716, Table S12). As shown previously (84), males avoid leg vibration when the risk of interception is high (e.g., at high density). Hence, notwithstanding the lesser importance of resource-defense and body size for mating success in male-biased populations, males did not adopt alternative mating tactics. This raises the question of whether selection might instead favor increased investment in post-mating processes.

### Fertilization success

Across all treatments, male mating success (MS; 48.9%) explained most of the variation in reproductive success, followed by fertilization success (FS; 14.3%) and female fecundity (Fec; 6.4%; Fig. S5, Table S16). This pattern was comparable across social contexts in general, but while mating success explained three times the variation of fertilization success under an even OSR (MS: 42.5%, FS: 14.4%, Fec: 9.2%), these variance components were more similar under a male-biased OSR (MS: 37.3%, FS: 25.4%, Fec: 7.2%) despite unchanged female mating rates (*z* = 0.01, *P* = 0.99). Fertilization success may thus gain in relative importance when more potential mates compete over each mating opportunity, hampering female monopolization.

Males that were a female’s last mate until the end of the experiment (*t =* 4.19, *P* < 0.001, *n* = 190, Table S16), and males that were able to delay remating for longer (“remating interval”, see *SI Methods; t* = 5.00, *P* < 0.001), had higher fertilization success (adjusted PCS, see *SI Methods*). Analogous to our findings for mating success, larger males only had higher fertilization success under an even OSR (*t* = 2.68, *P* = 0.008), but not under a male bias (*t* = 0.14, *P* = 0.88; OSR×PC1: *t* = 1.83, *P* = 0.069, *n* = 190, Table S16). Longer mating durations did not significantly contribute to fertilization success (*t* = 1.36, *P* = 0.18), but males mating more often had lower fertilization success per mating (*t* = −2.22, *P* = 0.027), suggesting strategic allocation (85) or sperm depletion (86).

Female remating intervals were not affected by density (*t* = 0.79, *P* = 0.43), but they were longer under even than male-biased OSRs (*t* = 1.90, *P* = 0.06, *n* = 190, Table S17). Copulation duration was positively related with remating intervals (*t* = 1.81, *P* = 0.073), but did not vary with OSR or density (both |*t*| < 1.41, *P* > 0.17, *n* = 710). These were unexpected to the extent that male *Drosophila* have been shown to transfer more of the seminal fluid proteins (SFPs) known to suppress female remating when they perceive a high sperm competition risk (e.g., 87), and this allocation may covary with copulation duration (88). Our results, however, support previous findings suggesting a limited role for the control of remating suppression through SFPs in this (72) and other species with high mating rates (89). If remating intervals are under female control, females might delay remating until a favored male is found (90, 91). Some support for this idea comes from the fact that large males were more likely to be the last male overall (i.e., until the end of the experiment; *t* = 2.25, *P* = 0.026, *n* = 211), but size did not affect remating intervals up to that point (*t* = 0.05, *P* = 0.95).

In sum, our results indicate that intensifying competition does not augment post-mating investment by males. Additionally, post-mating selection does not seem to counteract the lack of pre-mating selection on ostensibly beneficial male traits. Instead, post-mating processes seem to reinforce the patterns of pre-mating selection.

### Mating system dynamics

To synthesize the differences in the mating-system dynamics of *D. prolongata* across different contexts, we built structural equation models. We provide separate visualizations for each density level in Fig. S6 (Tables S22-S24), but Fig. 4 presents a simplified overview of the dynamics under even and male-biased OSRs, respectively. One of the most striking observations was the absence of many positive (green) or negative (red) effects in the male-biased scenario (Fig. 4b, Table S21) compared to that under even OSR (Fig. 4a, Table S20). This underlines how both pre- and post-mating sexual selection vary across social contexts. For example, variation in body size, a primary determinant of both pre- and post-mating success under even OSR, seemed inconsequential under male-biased OSRs. Although we documented a larger skew in reproductive success and a higher potential for sexual selection in male-biased populations, we were unable to pinpoint the specific traits under selection. Altered ecological or demographic factors can result in plastic changes in mating systems and ultimately individual strategies (7, 37, 45, 64). When the number of rivals increases (e.g., male-biased OSR) or the encounter rate with mates declines (e.g., low density), males might transition from territorial contests to an alternative mating tactic, potentially a form of scramble competition (49, 50, 79). Scramble competition polygyny is a common mating system in invertebrates and can manifest in diverse forms (79). At low density, females may be dispersed, generating selection on the ability to locate receptive females (79). In another type of scramble competition, individuals of both sexes cluster around resources that cannot be economically monopolized (e.g., if resource is widely distributed or number of competitors is high). In this form of scramble competition, it can be a race between competitors to locate a receptive female in a dense aggregation, possibly combined with coercion or courtship when the female is reached (79). In such scenarios, traits like speed, agility, or the ability to locate receptive females could be under selection (79).

**Figure 4:**
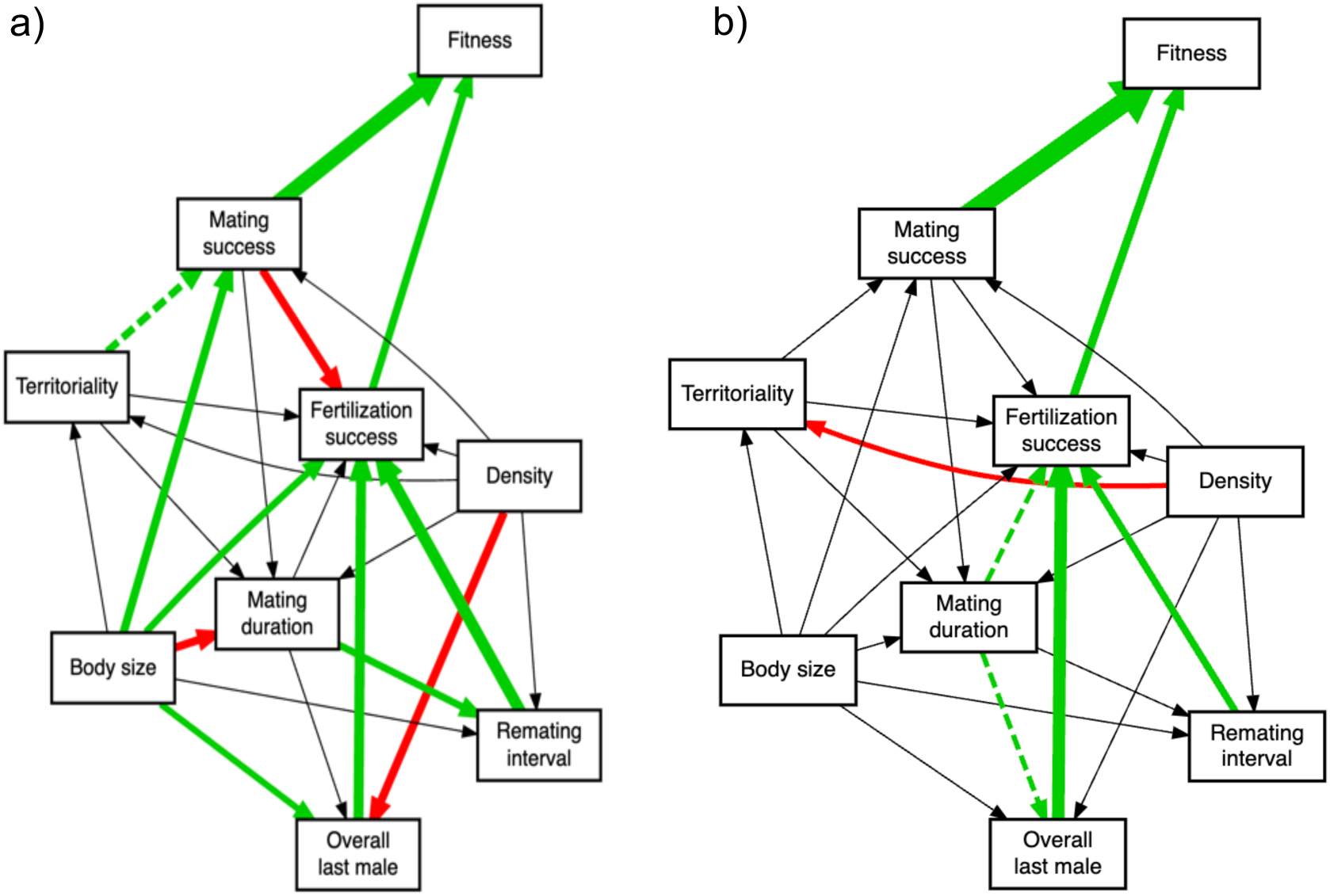
Visualized structural equation model of male reproductive dynamics under (a) an even OSR. Green and red indicate positive and negative relationships, respectively. Black lines indicate neutral relationships. Dashed lines (red or green) indicate trends (*P* < 0.1).

While not systematically quantifying them in this study, we did frequently observe multi-male clusters (‘love knots’ (80)) when a receptive female had been located, often followed by mating. Consequently, morphological traits linked to speed may also be under selection like their behavioral counterparts (79). Incidentally, of all morphological traits assessed here, the only one exhibiting significant positive selection under male-biased OSRs was wing length, a trait often thought to be selected under scramble competition due to its link to speed (79). Similarly, small body size has been proposed to be advantageous in scramble competition (2, 79, 92), with selection on it documented in several arthropods (e.g., 93, 94), including multi-male scramblers (80). Here, we did not measure the behavioral traits potentially under selection and found no significant negative selection on body size under male-biased OSR or low density. However, we have unpublished data after experimentally evolving replicated lines under different OSRs for 35 generations, which indicate a reduction in male body size under male-biased relative to even OSR (*t* = −6.58, *n* = 681, *P* < 0.001). In combination with the results of this study, this divergence may indicate a shift to a mating system broadly described as scramble competition, or certainly away from territorial contest competition. It is important to note, however, that contest and scramble competition are not necessarily separate categories but rather “extremes on a continuum”(79, p. 247). Considering the pronounced male-biased sexual size dimorphism in *D. prolongata,* which is unique among Drosophilids (95) and uncommon for species reliant on scramble competition (92, 96), scramble competition might play a minor role in this species, triggered primarily under relatively extreme (and potentially unnatural) circumstances. Whether a shift toward scramble competition would occur at low densities or under a male-biased OSR remains unclear, as our treatments were not extreme enough or the sample size too low to conclusively detect it.

## Conclusion

Our findings broaden our current understanding of how mating systems respond to complex socio-ecological pressures. The plastic mating strategies observed in *D. prolongata* challenge the notion of static classifications in favor of a dynamic framework, likely beyond this species. This dynamism emphasizes the importance of studying selection across ecological contexts, such as varying OSRs, population densities or resource distributions to understand the intricate interplay between environmental change, population demographics, and the evolutionary process. Incorporating these factors expands our perspective beyond single-environment selection metrics, with important implications. For example, traits like body size may be associated with dominance in contest competition and so assumed to be under directional selection, but this does not mean that the same traits are also advantageous in other contexts (e.g., dynamic scramble competition (79)) or where post-mating selection becomes more pronounced. Our study also highlights that integrating multiple indices of sexual selection intensity and complementing these with measures of selection on phenotypic traits offers a far more comprehensive picture of how sexual selection shapes phenotypes. Finally, by tracking individuals over multiple mating events we were able to quantify how post-reinforces pre-mating selection and how the targets of selection change across sequential matings. Our integrated approach holds the potential to advance our understanding of evolutionary adaptation and pave the way for the development of more robust predictions about how populations will respond to future environmental challenges.

## Materials and Methods

### Study organism

The *D. prolongata* flies used in this study were descendants from 45 isofemale lines, originally collected in the Sa Pa region in Vietnam (66) maintained in the lab for approx. 30 generations at 20±1°C and 60% relative humidity, on a 14:10 light:dark cycle. To avoid potential inbreeding effects and maximize genetic diversity within each experimental unit, virgin males and females were crossed between independent lines, with varying heterozygous genotypes between units. To ensure virginity of experimental individuals, flies were collected by aspiration on the day of eclosion before being kept in sex-specific vials (groups of 5 males or 10 females of the same parental genotypes) on standard fly food (see *SI Methods*) until sexually mature at 5-7 days post-eclosion. For identification, the thorax of each experimental fly was marked with a dot of acrylic paint (Kreul El Greco Acrylic colors) in a unique color within sex and experimental unit, three days before the start of the experiment. Thereafter, flies were housed individually in 20mL vials with approx. 2mL of fly food.

### Experimental units and procedure

Each experimental arena (see *SI Methods*) consisted of a Petri dish (ø 9cm) with small ventilation holes in the lid. Petri dishes were filled with orange juice-agar medium (see *SI Methods*) around a central Petri dish (ø 3.5cm) that was filled with standard fly food and covered with yeast paste. This central yeast patch served a resource-dense area (RDA), whilst the peripheral agar medium was considered resource-poor, with small grains of dry yeast sprinkled at a low density (Fig. S7). Each experimental unit (group of flies) was housed in one of these Petri dishes, varying in their operational sex ratio (OSR; 1:1 or 3:1 M:F) and population density (low [12 flies], medium [16], high [20]). Each group composition was replicated five times.

Flies were introduced into the plates by aspiration according to the density and sex ratio of their treatment. There was neither an overlap of isoline crosses within a plate nor repeated color within either sex of a plate. Each plate was placed under a Raspberry Pi 3 model B attached to a camera module (V2) in a climate chamber at 20°C and 60% humidity with a 14:10 light:dark cycle, and fans near the plates prevented condensation of the lids. Units were photographed from above every two seconds during the light period across five days. On days two to five, groups of flies were moved to fresh plates within the first hour of light. All eggs and larvae of the previous plate were transferred to a fresh plate containing standard fly food and left to develop for 3-4 days before being collected in individual Eppendorf tubes and stored at −20°C for later molecular parentage analysis. On the morning after the fifth night, all experimental flies were removed and anesthetized on ice before separately storing their heads and bodies in Eppendorf tubes at −20°C for later genotyping and morphometry, respectively.

### Morphological measurements

For each male and female, thorax length was measured, using the reticular eyepiece of a Leica M60 stereoscope. In males, the left foreleg and wing were removed using forceps and mounted on a microscope slide in a drop of Euparal underneath a coverslip. These body parts were then photographed at 25x magnification using a sCMOS Microscope camera K5 on a Leica M165 FC stereoscope to measure wing length and total foreleg length using Fiji (97) (details in *SI Methods*). Male thorax, wing and foreleg length were further combined in a principal component analysis, with the first principal component (PC1) serving as a proxy of overall body size (see *SI Methods*).

### Collection of behavioral data

The images were visually screened for matings. For each mating, we recorded the male and female ID’s, along with the start and end times to calculate mating durations and remating intervals. We further noted whether copulations started in the RDA, whether they were attained by an alternative reproductive tactic (“interception”, (84)), aggression or courtship, and whether they were preceded by leg vibration (see *SI Methods* for details). In addition to quantifying matings and accompanying behaviors, we recorded every 10 minutes which individuals were present in the RDA (i.e. at least their head or the thorax and abdomen were in the RDA).

### Parentage analysis

The parentage analysis was based on six microsatellite loci that we previously determined to be highly variable among isolines (details in *SI Methods*). The larvae or heads of parents were suspended in KAWA buffer (10 mM Tris HCl, 1 mM EDTA, 0.5% Tween 20, 50 µg/ml Proteinase K, pH 8.0) and sent to Ecogenics (Microsynth) for DNA extraction, PCR amplification and sequencing. Approximately 50% of all 10,667 larvae were selected at random from each plate and day for genotyping. Parentage assignment was done using CERVUS 3.0.7 (98), which determines the most likely parent-offspring trio by maximum likelihood exclusion based on LOD (logarithm of the odds) scores and their probabilities by simulations on the allele frequencies in the population. Parent-offspring trios were accepted as valid when at least three loci were compared (excluded when more than three were missing), and when these were assigned with a likelihood of 95% (“strict”) or 80% (“relaxed”). As all potential parents were known and combinations could be cross-checked with observed matings, trios with 80% likelihood were included.

### Statistical analyses

All statistical analyses were conducted in R version 4.3. (99). Linear and generalized linear mixed models were conducted using the package *lme4* (100) with experimental units as a random effect. Where the analysis was done on the level of matings instead of individuals, the unique male and female ID were added as random effects. Negative binomial model (*glmmTMB* (101)) or observation level random effects (102) were used to control for overdispersion in Poisson or binomial models, respectively. We bootstrapped confidence intervals (n = 1000) for the opportunity for sexual selection and for the variance partitioning using the package *sampling* (103). The morphological measurements for the selection gradients were scaled within each experimental unit, and relative reproductive success within experimental units was used as a response variable. We used the package *piecewiseSEM* (104) for piecewise structural equation modeling and *diagrammeR* (105) to visualize the structural equation models. A comprehensive description of all statistical methods can be found in the *SI Methods*.

## Acknowledgments

We thank Florin Allmendinger, Sina Lerch, Monica Anderson Berdal, Sonja Sbilordo and Jeannine Roy for their support in the lab, and Scott Pitnick for valuable comments. This project was supported by the Swiss National Science Foundation (grants 310030_197651 and PP00P3_202662 to S.L.) and by the University of Zurich (FK-20-088 to A.D.N.).

## Notes

### Competing Interest Statement

The authors have declared no competing interest.

### Summary of Updates

The manuscript has been revised for clarity, and some analyses and supplementary figures have been added.

