## Supplementary information for "Socio-ecological context modulates significance of territorial contest competition in *Drosophila prolongata*"

**Supplementary Information for:**
**"Socio-ecological context modulates significance of territorial contest competition in *Drosophila prolongata*"**

Alessio N. De Nardo*, Broti Biswas, Jhoniel Perdigón Ferreira, Abhishek Meena, Stefan Lüpold

### Supplementary Results

#### Figures

**Figure S1:** The violin plot shows the distributions of relative mating success for males in the even and male-biased OSR, respectively. The numbers next to the lines on the left side, indicate the percentiles and the lines show how they differ between the OSRs.

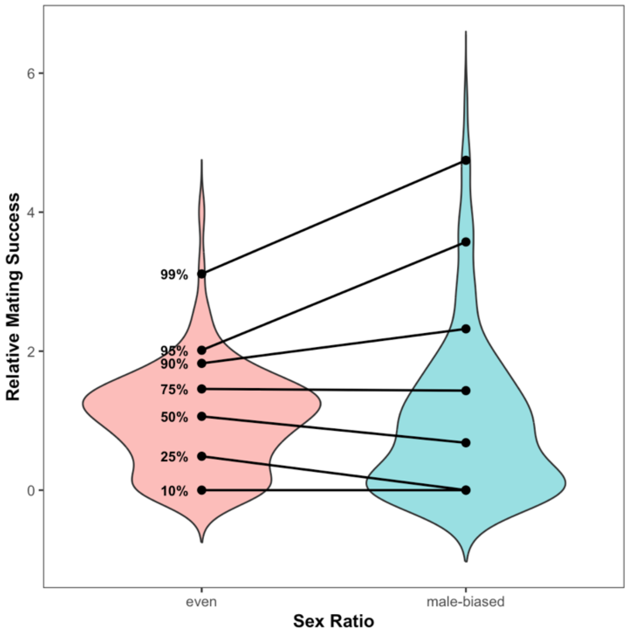

**Figure S2:** The violin plot shows the distributions of relative mating success for males in the low, medium and high density treatments, respectively. The numbers next to the lines on the left side, indicate the percentiles and the lines show how they differ between the densities.

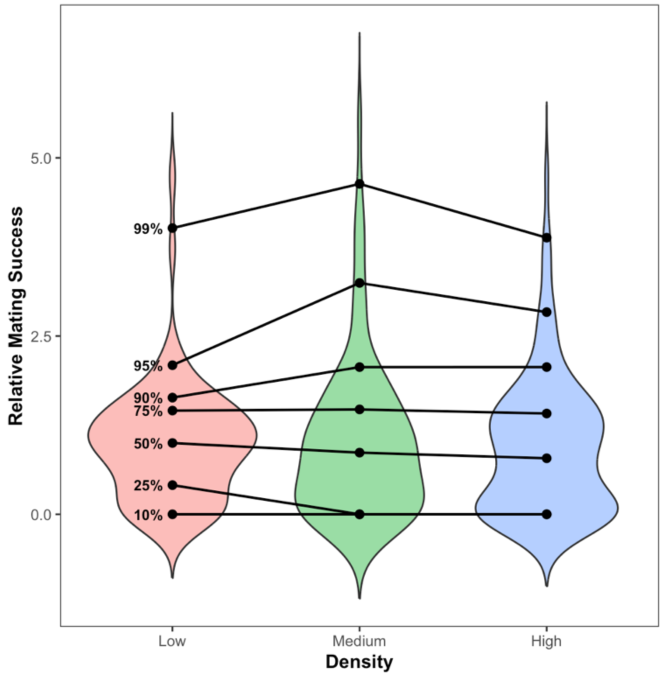


**Figure S3:** The opportunity for sexual selection (mean ± bootstrapped 95% confidence intervals [$n$ = 1000]) in different socio-ecological contexts: even (E) or male-biased (MB) OSR and low, medium, or high density. $I_s$ of females and males are visualized in red and blue, respectively. Means with 95% confidence intervals. This figure corresponds to Table S4.

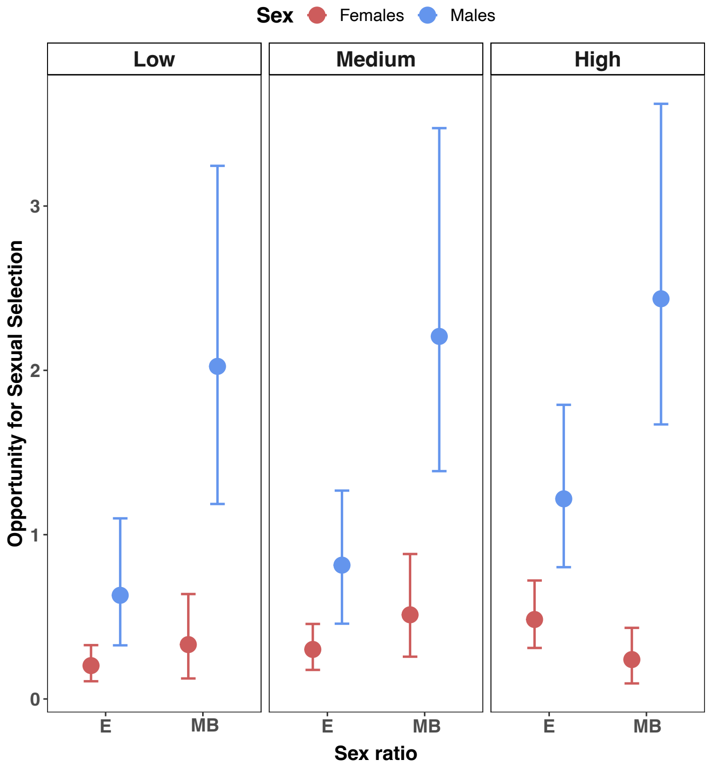

**Figure S4:** Percentage of time observed on food patch (mean ± 95% confidence interval) depending on the sex and day. The figure corresponds to Table S8.

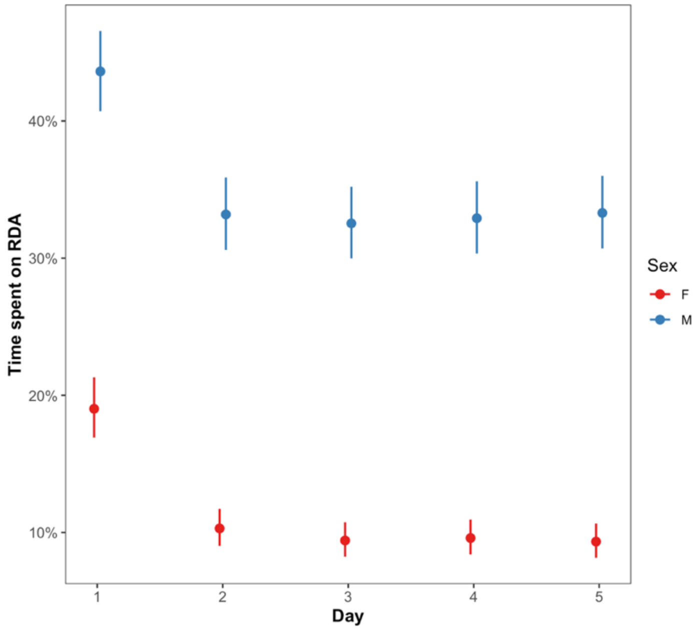


**Figure S5:** Variance partitioning of reproductive success (mean ± bootstrapped 95% confidence intervals [*n* = 1000]) for mating success (MS), fertilization success (FS), and average fecundity of the females (Fec). This figure corresponds to Table S15.

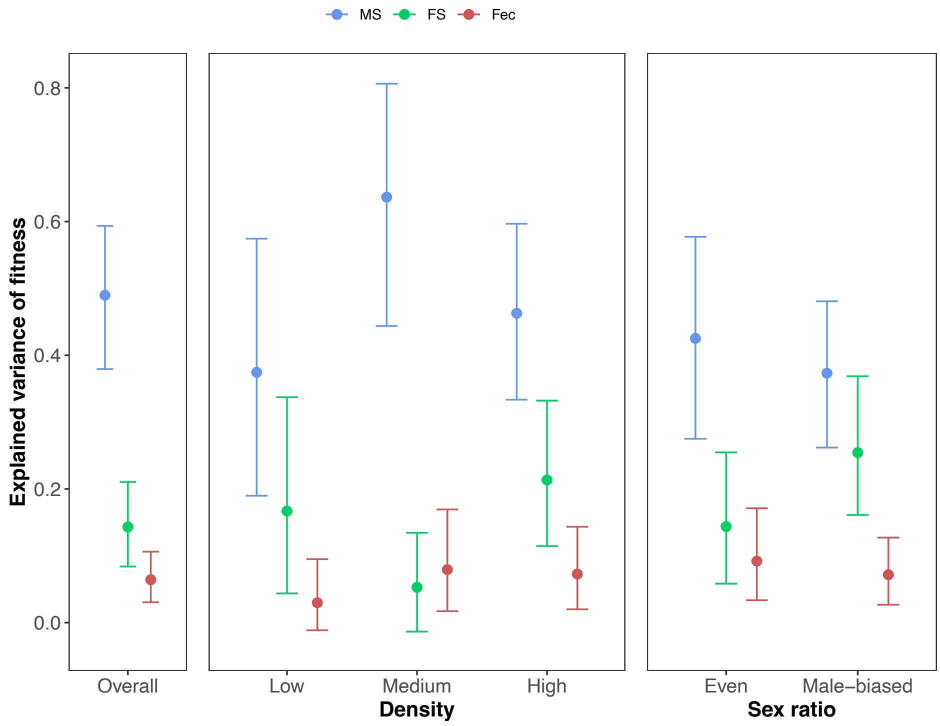

**Figure S6:** Visualization of structural equation models of male reproductive dynamics under low, medium, and high density (left to right). Green and red indicate positive and negative relationships, respectively. Black lines indicate neutral relationships. Dashed lines (red or green) indicate trends ($P < 0.1$). The figures correspond to Tables S22-24, respectively.

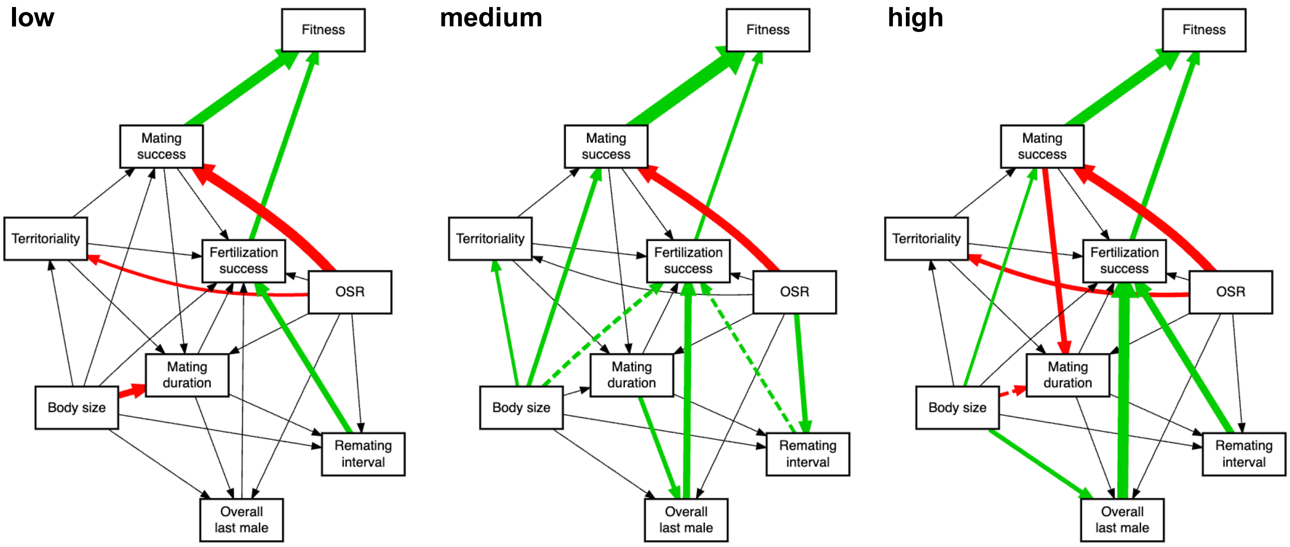


### Tables

**Table S1:** ANOVA table for Bateman gradients in males (GLMM; *n* = 193). Bold *P*-values are significant ($P < 0.05$) and italic *P*-values indicate a trend ($P < 0.1$).

| | $\chi^2$ | df | *P* |
|---|---|---|---|
| Mates | 63.65 | 1 | **< 0.001** |
| OSR | 1.74 | 1 | 0.19 |
| Density | 1.72 | 2 | 0.42 |
| Mates:OSR | 2.8 | 1 | *0.09* |
| Mates:Density | 2.59 | 2 | 0.27 |
| OSR:Density | 5.93 | 2 | *0.051* |
| Mates:OSR:Density | 0.008 | 2 | 0.99 |

**Table S2:** ANOVA table for Bateman gradients in females (GLMM; *n* = 175). Bold *P*-values are significant ($P < 0.05$) and italic *P*-values indicate a trend ($P < 0.1$).

| | $\chi^2$ | df | *P* |
|---|---|---|---|
| Number of mates (Mates) | 24.08 | 1 | **< 0.001** |
| OSR | 0.06 | 1 | 0.80 |
| Density | 0.86 | 2 | 0.65 |
| Mates:OSR | 1.55 | 1 | 0.22 |
| Mates:Density | 1.64 | 2 | 0.44 |
| OSR:Density | 0.16 | 2 | 0.92 |
| Mates:OSR:Density | 0.80 | 2 | 0.67 |

**Table S3a:** ANOVA table for Bateman gradients across the sexes (negative binomial; *n* = 368). OSR and density combined to "treatment". Corresponds to Figure 1 in the main text. Bold *P*-values are significant ($P < 0.05$) and italic *P*-values indicate a trend ($P < 0.1$).

| | $\chi^2$ | df | *P* |
|---|---|---|---|
| Number of mates (Mates) | 81.64 | 1 | **< 0.001** |
| Treatment | 3.39 | 5 | 0.63 |
| Sex | 1.57 | 1 | 0.21 |
| Mates:Treatment | 3.45 | 5 | 0.63 |
| Mates:Sex | 7.42 | 1 | **0.0064** |
| Treatment:Sex | 4.84 | 5 | 0.44 |
| Mates:Treatment:Sex | 6.02 | 5 | 0.30 |

**Table S3b:** ANOVA table for Bateman gradients across the sexes (negative binomial; *n* = 368). OSR and density separate. Bold *P*-values are significant ($P < 0.05$) and italic *P*-values indicate a trend ($P < 0.1$).

| | $\chi^2$ | df | *P* |
|---|---|---|---|
| Number of mates (Mates) | 81.64 | 1 | **< 0.001** |
| OSR | 0.25 | 1 | 0.62 |
| Density | 1.79 | 2 | 0.41 |
| Sex | 1.57 | 1 | 0.21 |
| Mates:OSR | 0.21 | 1 | 0.64 |
| Mates:Density | 3.26 | 2 | 0.20 |
| OSR:Density | 1.71 | 2 | 0.43 |
| Mates:Sex | 7.42 | 1 | **0.006** |
| OSR:Sex | 1.20 | 1 | 0.27 |
| Density:Sex | 0.51 | 2 | 0.77 |
| Mates:OSR:Density | 0.23 | 2 | 0.89 |
| Mates:OSR:Sex | 5.60 | 1 | **0.018** |
| Mates:Density:Sex | 0.37 | 2 | 0.83 |
| OSR:Density:Sex | 3.22 | 2 | 0.20 |
| Mates:OSR:Density:Sex | 0.32 | 2 | 0.85 |

**Table S4:** Summary table for the opportunity for sexual selection, $I_S$, with means and bootstrapped 95% confidence limits (*n* = 1000) for each group (divided by sex, OSR, density), and the male:female ratio of the $I_S$. Bold *P*-values are significant ($P < 0.05$) and italic *P*-values indicate a trend ($P < 0.1$). The results are visualized in Figure S1.

| Sex | OSR | Density | $I_s$ with 95% confidence limits | Ratio $I_s$ Male:Female |
|---|---|---|---|---|
| F | Even | Low | 0.20 [0.11, 0.30] | 3.1 |
| M | Even | Low | 0.62 [0.31, 1.16] | |
| F | Male-biased | Low | 0.31 [0.13, 0.64] | 6.6 |
| M | Male-biased | Low | 2.05 [1.17, 3.41] | |
| F | Even | Medium | 0.29 [0.18, 0.42] | 2.7 |
| M | Even | Medium | 0.78 [0.44, 1.23] | |
| F | Male-biased | Medium | 0.53 [0.27, 0.90] | 4.3 |
| M | Male-biased | Medium | 2.28 [1.43, 3.58] | |
| F | Even | High | 0.49 [0.31, 0.70] | 2.5 |
| M | Even | High | 1.21 [0.81, 1.73] | |
| F | Male-biased | High | 0.21 [0.11, 0.34] | 11.2 |
| M | Male-biased | High | 2.36 [1.68, 3.70] | |

**Table S5:** Slopes, Standard errors (SE), 95% confidence limits (CL) and *P*-values for selection gradients for the different OSRs. Contrasts comparing gradients pairwise between different OSRs. Bold *P*-values are significant ($P < 0.05$) and italic *P*-values indicate a trend ($P < 0.1$). This table corresponds to the right side of Figure 2 in the main text.

| | | | | | | | Contrasts | |
|---|---|---|---|---|---|---|---|---|
| **Trait** | **OSR** | **Slope** | **SE** | **Lower CL** | **Upper CL** | ***P*-value slope** | ***z*** | ***P*** |
| Thorax length | Even | 0.4 | 0.08 | 0.24 | 0.57 | **<0.001** | 2.55 | **0.011** |
| | Male-biased | 0.03 | 0.12 | -0.19 | 0.27 | 0.78 | | |
| Foreleg length | Even | 0.36 | 0.09 | 0.18 | 0.53 | **<0.001** | 1.52 | 0.13 |
| | Male-biased | 0.13 | 0.12 | -0.09 | 0.36 | 0.25 | | |
| Wing length | Even | 0.41 | 0.09 | 0.24 | 0.58 | **<0.001** | 1.07 | 0.28 |
| | Male-biased | 0.25 | 0.12 | 0.02 | 0.48 | *0.03* | | |
| PC1 | Even | 0.4 | 0.09 | 0.23 | 0.57 | **<0.001** | 1.79 | *0.07* |
| | Male-biased | 0.14 | 0.12 | -0.09 | 0.37 | 0.24 | | |

**Table S6:** Slopes, 95% confidence limits (CL), Standard errors (SE), and *P*-values for selection gradients for the different densities. Contrasts comparing gradients pairwise between different densities. Bold *P*-values are significant ($P < 0.05$) and italic *P*-values indicate a trend ($P < 0.1$). This table corresponds to the left side of Figure 2 in the main text.

| | | | | | | | Contrasts with High | | Contrasts with Medium | | Contrasts with Low | |
|---|---|---|---|---|---|---|---|---|---|---|---|---|
| **Trait** | **Density** | **Slope** | **Lower CL** | **Upper CL** | **SE** | ***P*-value slope** | ***z*** | ***P*** | ***z*** | ***P*** | ***z*** | ***P*** |
| Thorax length | Low | -0.10 | -0.38 | 0.18 | 0.14 | 0.48 | 2.14 | **0.03** | 1.56 | 0.11 | - | - |
| | Medium | 0.2 | -0.06 | 0.47 | 0.13 | 0.13 | 0.59 | 0.55 | - | - | 1.56 | 0.11 |
| | High | 0.31 | 0.05 | 0.57 | 0.13 | **0.02** | - | - | 0.59 | 0.55 | 2.14 | **0.03** |
| Foreleg length | Low | -0.04 | -0.31 | 0.24 | 0.14 | 0.8 | 1.94 | *0.052* | 1.5 | 0.13 | - | - |
| | Medium | 0.25 | -0.01 | 0.51 | 0.13 | *0.06* | 0.46 | 0.64 | - | - | 1.5 | 0.13 |
| | High | 0.34 | 0.08 | 0.60 | 0.13 | **0.012** | - | - | 0.46 | 0.64 | 1.94 | *0.052* |
| Wing length | Low | -0.04 | -0.33 | 0.24 | 014 | 0.75 | 2.7 | **0.007** | 2.09 | **0.004** | - | - |
| | Medium | 0.36 | 0.10 | 0.62 | 0.13 | **0.01** | 0.64 | 0.52 | - | - | 2.09 | **0.004** |
| | High | 0.48 | 0.22 | 0.73 | 0.13 | **< 0.001** | - | - | 0.64 | 0.52 | 2.7 | **0.007** |
| PC1 | Low | -0.08 | -0.36 | 0.20 | 0.14 | 0.56 | 2.52 | **0.012** | 1.76 | *0.079* | - | - |
| | Medium | 0.26 | -0.004 | 0.52 | 0.13 | *0.053* | 0.78 | 0.43 | - | - | 1.76 | *0.079* |
| | High | 0.41 | 0.14 | 0.67 | 0.13 | **0.003** | - | - | 0.78 | 0.43 | 2.52 | **0.012** |

**Table S7:** ANOVA table for GLMM on the proportion of time spent on food overall (n = 493). Bold *P*-values are significant (*P* < 0.05) and italic *P*-values indicate a trend (*P* < 0.1).

| | $\chi^2$ | *df* | *P* |
|---|---|---|---|
| Sex | 671.6 | 1 | **< 0.001** |
| OSR | 20.9 | 1 | **< 0.001** |
| Density | 22.0 | 1 | **< 0.001** |

**Table S8:** ANOVA table for GLMM on the proportion of time spent on the RDA (per day) depending on sex and the day (n = 2379). Bold *P*-values are significant (*P* < 0.05) and italic *P*-values indicate a trend (*P* < 0.1). This table corresponds to Figure S2.

| | $\chi^2$ | *df* | *P* |
|---|---|---|---|
| Day (factor) | 313.7 | 4 | **< 0.001** |
| Sex | 658.6 | 1 | **< 0.001** |
| Day:Sex | 26.3 | 4 | **< 0.001** |

**Table S9:** ANOVA table for GLMM on the proportion of time spent on the RDA (per day) by males depending on PC1 and the day [first day or later] (n =1445). Bold *P*-values are significant (*P* < 0.05) and italic *P*-values indicate a trend (*P* < 0.1).

| | $\chi^2$ | *df* | *P* |
|---|---|---|---|
| Day (factor) | 132.9 | 1 | **< 0.001** |
| PC1 | 1.74 | 1 | 0.19 |
| PC1:Day | 5.05 | 1 | **0.025** |

**Table S10a:** Summary table for LMM on mating success depending on the OSR, density, PC1, and territoriality [even OSR as baseline] (*n* = 290). Bold *P*-values are significant (*P* < 0.05) and italic *P*-values indicate a trend (*P* < 0.1).

| | ***Estimate*** | ***SE*** | ***t*** | ***P*** |
|---|---|---|---|---|
| Intercept (Even OSR) | 1.34 | 0.07 | 18.1 | **< 0.001** |
| OSR (MB) | -0.69 | 0.08 | -9.0 | **< 0.001** |
| PC1 | 0.23 | 0.06 | 3.7 | **< 0.001** |
| Territoriality | 0.15 | 0.06 | 2.4 | **0.019** |
| Density (Low) | 0.25 | 0.09 | 2.7 | **0.007** |
| Density (Medium) | -0.02 | 0.09 | -0.3 | 0.80 |
| OSR (MB):PC1 | -0.17 | 0.08 | -2.2 | **0.03** |
| OSR (MB):Territoriality | -0.14 | 0.08 | -1.7 | *0.09* |

**Table S10b:** Summary table for LMM on mating success depending on the OSR, density, PC1, and territoriality [male-biased OSR as baseline] (*n* = 290). Bold *P*-values are significant (*P* < 0.05) and italic *P*-values indicate a trend (*P* < 0.1).

| | ***Estimate*** | ***SE*** | ***t*** | ***P*** |
|---|---|---|---|---|
| Intercept (MB OSR) | 1.34 | 0.07 | 18.1 | **< 0.001** |
| OSR (E) | 0.69 | 0.08 | 9.0 | **< 0.001** |
| PC1 | 0.06 | 0.05 | 1.1 | 0.26 |
| Territoriality | 0.01 | 0.05 | 0.2 | 0.81 |
| Density (Low) | 0.25 | 0.09 | 2.7 | **0.007** |
| Density (Medium) | -0.02 | 0.09 | -0.3 | 0.80 |
| OSR (E):PC1 | -0.17 | 0.08 | -2.2 | **0.03** |
| OSR (E):Territoriality | -0.14 | 0.08 | -1.7 | *0.09* |

**Table S11:** Summary table for the GLMM on whether leg vibration by the successful male was observed before a mating (*n* = 716). Bold *P*-values are significant (*P* < 0.05) and italic *P*-values indicate a trend (*P* < 0.1).

| | ***Estimate*** | ***SE*** | ***z*** | ***P*** |
|---|---|---|---|---|
| Intercept | -2.63 | 0.38 | -6.96 | **< 0.001** |
| Female mating status (virgin) | 0.68 | 0.27 | 2.56 | **0.01** |
| OSR (E) | 0.63 | 0.34 | 1.89 | *0.059* |
| PC1 | 0.30 | 0.11 | 2.89 | **0.004** |
| Density (numeric) | -0.32 | 0.19 | -1.70 | *0.089* |
| Mating on RDA (yes) | -0.48 | 0.26 | -1.81 | *0.07* |

**Table S12:** Summary table for the GLMM on whether a mating was gained by interception or not (*n* = 716). Bold *P*-values are significant (*P* < 0.05) and italic *P*-values indicate a trend (*P* < 0.1).

| | ***Estimate*** | ***SE*** | ***z*** | ***P*** |
|---|---|---|---|---|
| Intercept | -1.57 | 0.25 | -6.27 | **< 0.001** |
| Female mating status (virgin) | -0.16 | 0.23 | -0.73 | 0.46 |
| OSR (E) | -0.17 | 0.21 | -0.79 | 0.43 |
| PC1 | -0.13 | 0.06 | -2.03 | **0.042** |
| Density (numeric) | -0.14 | 0.12 | -1.13 | 0.26 |
| Mating on RDA (yes) | 0.52 | 0.22 | 2.31 | **0.021** |

**Table S13:** Summary table for the GLMM on whether a mating was preceded by an aggressive interaction (*n* = 716). Bold *P*-values are significant (*P* < 0.05) and italic *P*-values indicate a trend (*P* < 0.1).

| | ***Estimate*** | ***SE*** | ***z*** | ***P*** |
|---|---|---|---|---|
| Intercept | -1.09 | 0.20 | -5.45 | **< 0.001** |
| OSR (E) | -0.09 | 0.18 | -0.53 | 0.60 |
| PC1 | -0.05 | 0.05 | -0.89 | 0.37 |
| Density (numeric) | -0.17 | 0.10 | -1.60 | 0.11 |
| Mating on RDA (yes) | 0.48 | 0.19 | 2.54 | **0.011** |

**Table S14a:** Summary table for the GLMM on whether a mating was preceded by courtship by the successful male (*n* = 716; even OSR as baseline). Bold *P*-values are significant (*P* < 0.05) and italic *P*-values indicate a trend (*P* < 0.1).

| | ***Estimate*** | ***SE*** | ***z*** | ***P*** |
|---|---|---|---|---|
| Intercept | -1.13 | 0.19 | 5.78 | **< 0.001** |
| OSR (MB) | -0.03 | 0.19 | -0.18 | 0.85 |
| PC1 | 0.23 | 0.07 | 3.17 | **0.0014** |
| Density (numeric) | 0.14 | 0.11 | 1.24 | 0.22 |
| Mating on RDA (yes) | -0.65 | 0.20 | -3.3 | **< 0.001** |
| OSR (MB):PC1 | -0.36 | 0.12 | -2.93 | **0.0034** |

**Table S14b:** Summary table for the GLMM on whether a mating was preceded by courtship by the successful male (*n* = 716; male-biased OSR as the reference). Bold *P*-values are significant (*P* < 0.05) and italic *P*-values indicate a trend (*P* < 0.1).

| | ***Estimate*** | ***SE*** | ***z*** | ***P*** |
|---|---|---|---|---|
| Intercept | -1.10 | 0.21 | 5.19 | **< 0.001** |
| OSR (E) | 0.03 | 0.19 | 0.18 | 0.85 |
| PC1 | -0.12 | 0.09 | -1.27 | 0.20 |
| OSR (numeric) | 0.14 | 0.11 | 1.24 | 0.22 |
| Mating on RDA (yes) | -0.65 | 0.20 | -3.3 | **< 0.001** |
| OSR (E):PC1 | 0.36 | 0.12 | 2.93 | **0.0034** |

**Table S15:** Variance partitioning of reproductive success including bootstrapped 95% confidence intervals (*n* = 1000). This table corresponds to Figure S3.

| **Treatment** | | **Variance component** | **Mean explained variance** | **Lower 95% confidence limit** | **Upper 95% confidence limit** |
|---|---|---|---|---|---|
| Overall | | Mating success | 0.489 | 0.379 | 0.593 |
| | | Fertilization success | 0.143 | 0.084 | 0.210 |
| | | Fecundity | 0.064 | 0.030 | 0.106 |
| Density | Low | Mating success | 0.374 | 0.190 | 0.574 |
| | | Fertilization success | 0.167 | 0.044 | 0.337 |
| | | Fecundity | 0.029 | -0.011 | 0.095 |
| | Medium | Mating success | 0.637 | 0.444 | 0.806 |
| | | Fertilization success | 0.053 | -0.013 | 0.134 |
| | | Fecundity | 0.079 | 0.017 | 0.169 |
| | High | Mating success | 0.463 | 0.334 | 0.597 |
| | | Fertilization success | 0.213 | 0.115 | 0.332 |
| | | Fecundity | 0.073 | 0.020 | 0.143 |
| OSR | Even | Mating success | 0.425 | 0.275 | 0.577 |
| | | Fertilization success | 0.144 | 0.058 | 0.255 |
| | | Fecundity | 0.092 | 0.034 | 0.171 |
| | Biased | Mating success | 0.373 | 0.262 | 0.481 |
| | | Fertilization success | 0.254 | 0.161 | 0.369 |
| | | Fecundity | 0.072 | 0.027 | 0.127 |

**Table S16:** Summary table for the linear model on fertilization success ($n$ = 190). Bold $P$-values are significant ($P$ < 0.05) and italic $P$-values indicate a trend ($P$ < 0.1).

| | ***Estimate*** | ***SE*** | ***t*** | ***P*** |
|---|---|---|---|---|
| (Intercept) | -0.46 | 0.33 | -1.37 | 0.17 |
| PC1 | 0.04 | 0.01 | 2.68 | **0.008** |
| OSR (B) | -0.04 | 0.04 | -1.03 | 0.30 |
| Territoriality | -0.09 | 0.14 | -0.70 | 0.49 |
| Mean mating duration | 0.08 | 0.06 | 1.36 | 0.18 |
| Total matings | -0.02 | 0.01 | -2.22 | **0.027** |
| Mean remating interval (of mates) | 0.07 | 0.02 | 5.00 | **< 0.001** |
| overall last-male (proportion) | 0.32 | 0.08 | 4.19 | **< 0.001** |
| Density (numeric) | 0.01 | 0.02 | 0.21 | 0.83 |
| PC1:OSR (B) | -0.04 | 0.02 | -1.83 | *0.069* |

**Table S17:** Summary table for the linear model on the remating interval ($n$ = 190). Bold $P$-values are significant ($P$ < 0.05) and italic $P$-values indicate a trend ($P$ < 0.1).

| | *Estimate* | *SE* | *t* | *P* |
|---|---|---|---|---|
| Intercept | 3.54 | 1.59 | 2.23 | **0.03** |
| PC1 | 0.01 | 0.05 | 0.05 | 0.95 |
| overall last-male (proportion) | -0.08 | 0.37 | -0.23 | 0.82 |
| Mean mating duration | 0.50 | 0.28 | 1.81 | *0.073* |
| OSR (B) | -0.29 | 0.15 | -1.90 | *0.059* |
| Density (numeric) | 0.07 | 0.09 | 0.79 | 0.43 |

**Table S18:** Summary table for the linear model on the proportion of times the focal male was the last male overall ($n$ = 211). Bold $P$-values are significant ($P$ < 0.05) and italic $P$-values indicate a trend ($P$ < 0.1).

| | *Estimate* | *SE* | *t* | *P* |
|---|---|---|---|---|
| Intercept | 0.21 | 0.02 | 8.82 | **< 0.001** |
| PC1 | 0.02 | 0.01 | 1.98 | **0.048** |
| OSR (B) | - 0.02 | 0.03 | -0.62 | 0.54 |
| Mean mating duration | 0.02 | 0.02 | 1.37 | 0.17 |
| Density (numeric) | 0.04 | 0.02 | 1.88 | *0.062* |
| PC1:mean mating duration | -0.02 | 0.01 | -1.69 | *0.092* |

**Table S19:** Summary table for the linear model on mating duration (on the mating level; *n* = 710). Bold *P*-values are significant (*P* < 0.05) and italic *P*-values indicate a trend (*P* < 0.1).

| | *Estimate* | *SE* | *t* | *P* |
|---|---|---|---|---|
| Intercept | 332.9 | 11.10 | 30.01 | **< 0.001** |
| Density (numeric) | -11.78 | 8.39 | -1.41 | 0.17 |
| PC1 | -10.99 | 3.31 | -3.32 | **0.001** |
| OSR (E) | 4.47 | 14.23 | 0.31 | 0.76 |
| Prior matings (on that day) | 1.18 | 3.42 | 0.35 | 0.73 |
| Total matings (on that day) | -8.04 | 2.87 | -2.80 | **0.005** |
| Female mating status (virgin) | -23.52 | 8.55 | -2.75 | **0.006** |

**Table S20:** Summary table for the piecewise structural equation model for the even OSR. Bold *P*-values are significant (*P* < 0.05) and italic *P*-values indicate a trend (*P* < 0.1). This table corresponds to Figure 4a in the main text.

| ***Response*** | ***Predictor*** | **Estimate** | ***SE*** | ***df*** | ***t*** | ***P*** | ***Standardized estimate*** |
|---|---|---|---|---|---|---|---|
| Fitness | Mating success | 8.58 | 0.82 | 242.8 | 10.50 | **< 0.001** | **0.52** |
| Fitness | Fertilization success | 56.56 | 8.69 | 232.9 | 6.51 | **< 0.001** | **0.27** |
| Mating success | Body size | 0.49 | 0.17 | 69.8 | 2.87 | **0.005** | **0.29** |
| Mating success | Territoriality | 3.72 | 2.21 | 110.3 | 1.69 | *0.09* | *0.25* |
| Mating success | Density | -0.47 | 0.38 | 12.7 | -1.24 | 0.24 | -0.13 |
| Territoriality | Body size | 0.003 | 0.007 | 114 | 0.38 | 0.71 | 0.02 |
| Territoriality | Density | -0.023 | 0.015 | 114 | -1.55 | 0.12 | -0.09 |
| Fertilization success | Mating success | -0.022 | 0.007 | 86 | -3.04 | **0.003** | **-0.28** |
| Fertilization success | Territoriality | -0.25 | 0.15 | 86 | -1.64 | 0.11 | -0.22 |
| Fertilization success | Body size | 0.03 | 0.01 | 86 | 2.54 | **0.013** | **0.24** |
| Fertilization success | Density | -0.007 | 0.02 | 86 | -0.30 | 0.76 | -0.03 |
| Fertilization success | Remating interval | 0.09 | 0.02 | 86 | 4.85 | **< 0.001** | **0.40** |
| Fertilization success | Overall last male | 0.32 | 0.09 | 86 | 3.49 | **< 0.001** | **0.35** |
| Fertilization success | Mating duration | -0.15 | 0.09 | 86 | -1.59 | 0.12 | -0.14 |
| Remating interval | Body size | 0.03 | 0.07 | 90 | 0.46 | 0.65 | 0.05 |
| Remating interval | Density | 0.08 | 0.13 | 90 | 0.58 | 0.57 | 0.06 |
| Remating interval | Mating duration | 1.10 | 0.51 | 90 | 2.14 | **0.04** | **0.24** |
| Mating duration | Body size | -0.03 | 0.01 | 49.7 | -2.39 | **0.02** | **-0.26** |
| Mating duration | Density | -0.05 | 0.03 | 14.7 | -1.71 | 0.11 | -0.19 |
| Mating duration | Mating success | -0.008 | 0.01 | 87.1 | -0.95 | 0.35 | -0.10 |
| Mating duration | Territoriality | 0.11 | 0.18 | 89.6 | 0.62 | 0.53 | 0.10 |
| Overall last male | Body size | 0.03 | 0.02 | 93 | 2.02 | **0.046** | **0.22** |
| Overall last male | Density | 0.09 | 0.03 | 93 | 2.86 | **0.005** | **0.29** |
| Overall last male | Mating duration | 0.07 | 0.12 | 93 | 0.57 | 0.57 | 0.06 |

**Table S21:** Summary table for the piecewise structural equation model for the male-biased OSR. Bold *P*-values are significant (*P* < 0.05) and italic *P*-values indicate a trend (*P* < 0.1). This table corresponds to Figure 4b in the main text.

| ***Response*** | ***Predictor*** | **Estimate** | ***SE*** | ***df*** | ***t*** | ***P*** | ***Standardized estimate*** |
|---|---|---|---|---|---|---|---|
| Fitness | Mating success | 9.74 | 0.71 | 239.6 | 13.67 | **< 0.001** | **0.64** |
| Fitness | Fertilization success | 32.32 | 5.45 | 234.2 | 5.93 | **< 0.001** | **0.28** |
| Mating success | Body size | 0.09 | 0.08 | 109.8 | 1.22 | 0.22 | 0.07 |
| Mating success | Territoriality | 0.88 | 0.90 | 168.2 | 0.97 | 0.33 | 0.07 |
| Mating success | Density | -0.21 | 0.16 | 12.2 | -1.27 | 0.23 | -0.07 |
| Territoriality | Body size | 0.01 | 0.01 | 133.8 | 1.17 | 0.25 | 0.07 |
| Territoriality | Density | -0.04 | 0.01 | 15.1 | -2.50 | **0.02** | **-0.17** |
| Fertilization success | Mating success | -0.01 | 0.02 | 88 | -0.52 | 0.60 | -0.06 |
| Fertilization success | Territoriality | 0.05 | 0.19 | 88 | 0.25 | 0.81 | 0.03 |
| Fertilization success | Body size | 0.003 | 0.02 | 88 | 0.17 | 0.86 | 0.02 |
| Fertilization success | Density | 0.004 | 0.03 | 88 | 0.15 | 0.88 | 0.01 |
| Fertilization success | Remating interval | 0.07 | 0.02 | 88 | 2.78 | **0.006** | **0.24** |
| Fertilization success | Overall last male | 0.31 | 0.13 | 88 | 2.48 | **0.015** | **0.41** |
| Fertilization success | Mating duration | 0.15 | 0.08 | 88 | 1.99 | *0.05* | *0.17* |
| Remating interval | Body size | -0.01 | 0.07 | 92 | -0.10 | 0.92 | -0.01 |
| Remating interval | Density | 0.07 | 0.13 | 92 | 0.53 | 0.60 | 0.05 |
| Remating interval | Mating duration | 0.30 | 0.33 | 92 | 0.91 | 0.37 | 0.09 |
| Mating duration | Body size | -0.02 | 0.02 | 107.8 | -1.05 | 0.29 | -0.10 |
| Mating duration | Density | -0.02 | 0.05 | 12.7 | -0.32 | 0.75 | -0.04 |
| Mating duration | Mating success | -0.002 | 0.02 | 107.8 | -0.11 | 0.91 | -0.02 |
| Mating duration | Territoriality | 0.14 | 0.22 | 105.6 | 0.62 | 0.54 | 0.07 |
| Overall last male | Body size | 0.03 | 0.02 | 110 | 1.37 | 0.17 | 0.13 |
| Overall last male | Density | 0.01 | 0.04 | 110 | 0.32 | 0.75 | 0.03 |
| Overall last male | Mating duration | 0.21 | 0.11 | 110 | 1.89 | *0.06* | *0.17* |

**Table S22:** Summary table for the piecewise structural equation model for the low density. Bold *P*-values are significant (*P* < 0.05) and italic *P*-values indicate a trend (*P* < 0.1). This table corresponds to the left graph in Figure S4.

| ***Response*** | ***Predictor*** | **Estimate** | ***SE*** | ***df*** | ***t*** | ***P*** | ***Standardized estimate*** |
|---|---|---|---|---|---|---|---|
| Fitness | Mating success | 7.16 | 0.96 | 118.6 | 7.43 | **< 0.001** | **0.54** |
| Fitness | Fertilization success | 44.15 | 10.60 | 114.6 | 4.17 | **< 0.001** | **0.28** |
| Mating success | Body size | 0.04 | 0.21 | 68 | 0.19 | 0.85 | 0.02 |
| Mating success | Territoriality | 2.94 | 2.63 | 68 | 1.11 | 0.27 | 0.16 |
| Mating success | OSR | -1.56 | 0.31 | 68 | -5.11 | **< 0.001** | **-0.53** |
| Territoriality | Body size | -0.01 | 0.01 | 69 | -0.83 | 0.41 | -0.07 |
| Territoriality | OSR | -0.03 | 0.01 | 69 | -2.14 | **0.036** | **-0.18** |
| Fertilization success | Mating success | -0.01 | 0.02 | 47 | -0.96 | 0.34 | -0.17 |
| Fertilization success | Territoriality | -0.09 | 0.29 | 47 | -0.31 | 0.75 | -0.06 |
| Fertilization success | Body size | 0.01 | 0.02 | 47 | 0.32 | 0.75 | 0.05 |
| Fertilization success | OSR | -0.002 | 0.04 | 47 | -0.04 | 0.97 | -0.01 |
| Fertilization success | Remating interval | 0.08 | 0.03 | 47 | 2.41 | **0.019** | **0.31** |
| Fertilization success | Overall last male | 0.29 | 0.19 | 47 | 1.52 | 0.13 | 0.35 |
| Fertilization success | Mating duration | 0.12 | 0.12 | 47 | 0.97 | 0.33 | 0.14 |
| Remating interval | Body size | -0.09 | 0.11 | 48.2 | -0.79 | 0.43 | -0.13 |
| Remating interval | OSR | -0.14 | 0.15 | 7.9 | 0.92 | 0.38 | -0.15 |
| Remating interval | Mating duration | 0.12 | 0.51 | 48.9 | 0.23 | 0.82 | 0.04 |
| Mating duration | Body size | -0.07 | 0.03 | 53.8 | -2.55 | **0.01** | **-0.34** |
| Mating duration | OSR | -0.01 | 0.06 | 14.9 | -0.12 | 0.91 | -0.02 |
| Mating duration | Mating success | 0.001 | 0.02 | 54.6 | 0.09 | 0.93 | 0.01 |
| Mating duration | Territoriality | 0.08 | 0.30 | 51.7 | 0.27 | 0.79 | 0.05 |
| Overall last male | Body size | 0.03 | 0.03 | 56 | 0.87 | 0.39 | 0.13 |
| Overall last male | OSR | 0.03 | 0.04 | 56 | 0.83 | 0.41 | 0.11 |
| Overall last male | Mating duration | 0.01 | 0.15 | 56 | 0.05 | 0.96 | 0.01 |

**Table S23:** Summary table for the piecewise structural equation model for the medium density. Bold *P*-values are significant (*P* < 0.05) and italic *P*-values indicate a trend (*P* < 0.1). This table corresponds to the middle graph in Figure S4.

| ***Response*** | ***Predictor*** | **Estimate** | ***SE*** | ***df*** | ***t*** | ***P*** | ***Standardized estimate*** |
|---|---|---|---|---|---|---|---|
| Fitness | Mating success | 11.54 | 0.86 | 161.9 | 13.46 | **< 0.001** | **0.70** |
| Fitness | Fertilization success | 33.29 | 7.77 | 155.7 | 4.28 | **< 0.01** | **0.20** |
| Mating success | Body size | 0.40 | 0.16 | 94 | 2.55 | **0.01** | **0.24** |
| Mating success | Territoriality | 2.08 | 1.66 | 94 | 1.25 | 0.21 | 0.15 |
| Mating success | OSR | -1.17 | 0.25 | 94 | -4.68 | **< 0.001** | **-0.44** |
| Territoriality | Body size | 0.02 | 0.01 | 84.8 | 2.18 | **0.032** | **0.18** |
| Territoriality | OSR | -0.03 | 0.02 | 8.6 | -1.75 | 0.12 | -0.17 |
| Fertilization success | Mating success | -0.01 | 0.01 | 52 | -1.16 | 0.25 | -0.13 |
| Fertilization success | Territoriality | -0.30 | 0.21 | 52 | -1.48 | 0.15 | -0.22 |
| Fertilization success | Body size | 0.04 | 0.02 | 52 | 1.85 | *0.07* | *0.22* |
| Fertilization success | OSR | -0.05 | 0.03 | 52 | -1.58 | 0.12 | -0.19 |
| Fertilization success | Remating interval | 0.05 | 0.03 | 52 | 1.76 | *0.08* | *0.19* |
| Fertilization success | Overall last male | 0.28 | 0.13 | 52 | 2.20 | **0.03** | **0.37** |
| Fertilization success | Mating duration | 0.06 | 0.08 | 52 | 0.72 | 0.47 | 0.07 |
| Remating interval | Body size | -0.11 | 0.09 | 56 | -1.17 | 0.25 | -0.17 |
| Remating interval | OSR | -0.28 | 0.14 | 56 | -2.10 | **0.04** | **-0.28** |
| Remating interval | Mating duration | 0.55 | 0.40 | 56 | 1.38 | 0.17 | 0.18 |
| Mating duration | Body size | -0.01 | 0.03 | 62.3 | -0.43 | 0.67 | -0.06 |
| Mating duration | OSR | 0.01 | 0.05 | 13.0 | 0.25 | 0.81 | 0.04 |
| Mating duration | Mating success | 0.0001 | 0.02 | 60.2 | 0.01 | 0.99 | 0.001 |
| Mating duration | Territoriality | 0.46 | 0.30 | 62.6 | 1.51 | 0.14 | 0.26 |
| Overall last male | Body size | 0.01 | 0.03 | 62.4 | 0.49 | 0.62 | 0.06 |
| Overall last male | OSR | 0.03 | 0.04 | 6.3 | 0.61 | 0.56 | 0.08 |
| Overall last male | Mating duration | 0.25 | 0.12 | 62.8 | 2.01 | **0.048** | **0.24** |

**Table S24:** Summary table for the piecewise structural equation model for the high density. Bold *P*-values are significant (*P* < 0.05) and italic *P*-values indicate a trend (*P* < 0.1). This table corresponds to the right graph in Figure S4.

| ***Response*** | ***Predictor*** | **Estimate** | ***SE*** | ***df*** | ***t*** | ***P*** | ***Standardized estimate*** |
|---|---|---|---|---|---|---|---|
| Fitness | Mating success | 9.97 | 0.95 | 191.7 | 10.54 | **< 0.001** | **0.56** |
| Fitness | Fertilization success | 46.3 | 7.93 | 193.9 | 5.84 | **< 0.001** | **0.29** |
| Mating success | Body size | 0.23 | 0.11 | 89.2 | 2.07 | **0.04** | **0.17** |
| Mating success | Territoriality | 0.46 | 1.41 | 112.2 | 0.33 | 0.74 | 0.03 |
| Mating success | OSR | -1.16 | 0.26 | 6.72 | -4.55 | **0.003** | **-0.48** |
| Territoriality | Body size | -0.0002 | 0.007 | 117 | -0.03 | 0.98 | -0.002 |
| Territoriality | OSR | -0.04 | 0.01 | 117 | -3.47 | **< 0.001** | **-0.24** |
| Fertilization success | Mating success | -0.02 | 0.01 | 67 | -1.54 | 0.13 | -0.20 |
| Fertilization success | Territoriality | 0.09 | 0.20 | 67 | 0.46 | 0.65 | 0.06 |
| Fertilization success | Body size | 0.007 | 0.02 | 67 | 0.43 | 0.67 | 0.04 |
| Fertilization success | OSR | 0.004 | 0.03 | 67 | 0.11 | 0.91 | 0.01 |
| Fertilization success | Remating interval | 0.10 | 0.03 | 67 | 3.81 | **< 0.001** | **0.38** |
| Fertilization success | Overall last male | 0.47 | 0.13 | 67 | 3.61 | **< 0.001** | **0.57** |
| Fertilization success | Mating duration | 0.02 | 0.13 | 67 | 0.19 | 0.85 | 0.02 |
| Remating interval | Body size | 0.08 | 0.06 | 71 | 1.13 | 0.26 | 0.13 |
| Remating interval | OSR | -0.13 | 0.12 | 71 | -1.08 | 0.28 | -0.13 |
| Remating interval | Mating duration | 0.90 | 0.59 | 71 | 1.52 | 0.13 | 0.18 |
| Mating duration | Body size | -0.02 | 0.01 | 33.6 | -1.80 | *0.08* | *-0.19* |
| Mating duration | OSR | -0.02 | 0.03 | 9.1 | -0.58 | 0.58 | -0.08 |
| Mating duration | Mating success | -0.03 | 0.01 | 46.5 | -2.11 | **0.04** | **-0.30** |
| Mating duration | Territoriality | -0.12 | 0.18 | 76.2 | -0.69 | 0.49 | -0.11 |
| Overall last male | Body size | 0.04 | 0.02 | 79 | 2.24 | **0.03** | **0.24** |
| Overall last male | OSR | -0.05 | 0.03 | 79 | -1.37 | 0.17 | -0.15 |
| Overall last male | Mating duration | 0.23 | 0.17 | 79 | 1.36 | 0.18 | 0.15 |

**Supplementary Methods**

**Recipe for standard fly food.** The fly food on which the flies developed and were kept as adults until the start of the experiment consisted of the following ingredients per liter of food medium: 75 g glucose, 100 g fresh yeast, 55 g corn, 8 g agar, 10 g flour, and 15 mL Nipagin (antifungal).

**Experimental arena.** Each experimental arena was set up as shown in Figure S7, consisting of a petri dish was (ø 9cm) a smaller petri dish (ø 3.5cm) attached to the center. The periphery was filled with orange juice agar medium (600 mL water, 18g agar, 200mL orange juice, 20g sugar, 20mL Nipagin for a portion of approx. 850mL). The yeast paste covering the resource-dense area in the center was mixed from dry bakers' yeast and orange juice. Dry yeast was sprinkled on the orange juice agar in the periphery at approx. 5-6 kernels per $cm^2$ (counted in three randomly picked areas).

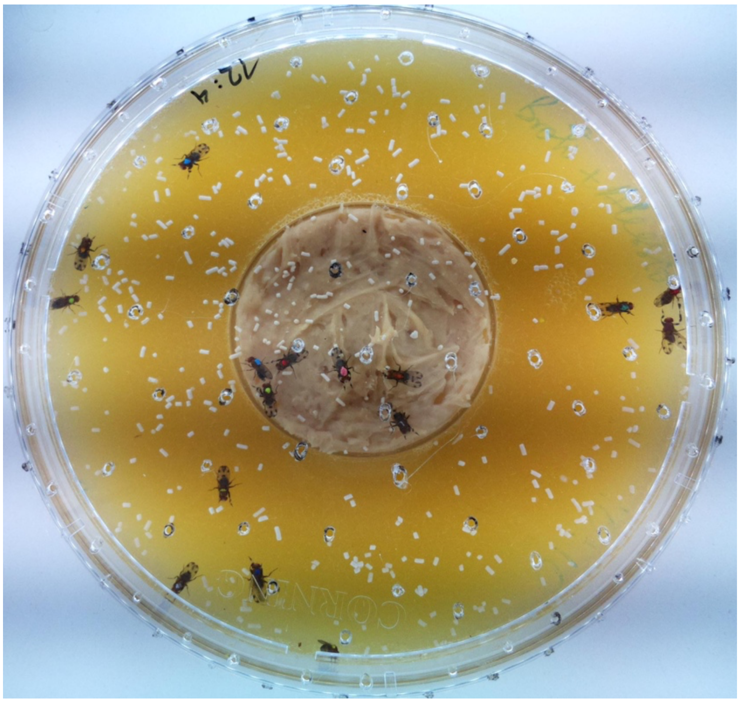

**Figure S7:** Example of experimental arena

**Morphological measurements.** The morphological traits used in this study (except for thorax length) were measured using Fiji (1). Figures S8 and S9 show how measurements were taken.

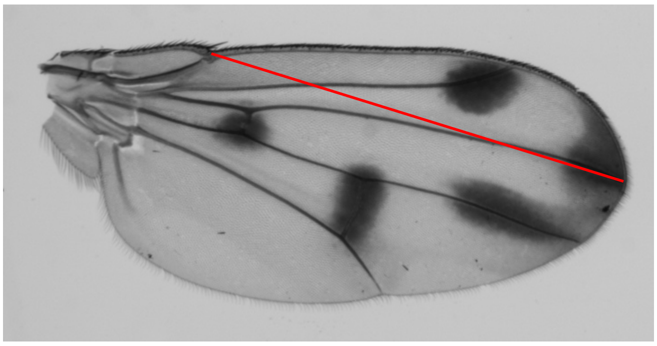

**Figure S8:** The line along which wing length was measure is shown in red and is clearly defined by identifiable landmarks on the wing.

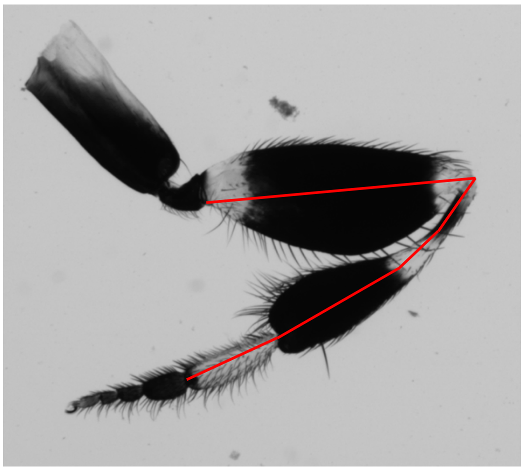

**Figure S9:** The line along which foreleg length was measured is shown in red and follows a segmented line through the femur, the tibia and the tarsus.

**Collection of behavioral data.** The images were visually screened for matings. Start and end times of matings, along with the male and female IDs were noted. For each mating, the male's behavior in the 30 seconds before the mating was assessed, noting whether courtship of the female, leg vibration or aggression toward another male occurred, whether the mating was attained by interception, and whether it started on the resource-dense area.

**Parentage analysis.** The parentage analysis was based on six microsatellite loci that we previously determined to be highly variable among isolines. Table S25 shows the primers that were used for the parentage analysis. The primer sequences will be made available on Genbank.

**Table S25:** Primers used for parentage analysis.

| Primer | Standard | Motif | Range of fragment length |
|---|---|---|---|
| G6 | AC | AC | 222-262 |
| G17 | AAC | TGT | 194-282 |
| G24 | AG | AG | 278-304 |
| G26 | AC | CA | 160-200 |
| G40 | AC | AC | 226-258 |
| G42 | ATC | ATC | 250-301 |

**Principal Component Analysis (PCA).** A PCA was conducted for PC1 to be used as a proxy of composite body size. The variables included were thorax, foreleg and wing length for all males with all these measurements available. The PCA was conducted in R version 4.3 using the function *prcomp* (2). A graphical representation of the PCA is shown in Figure S10.

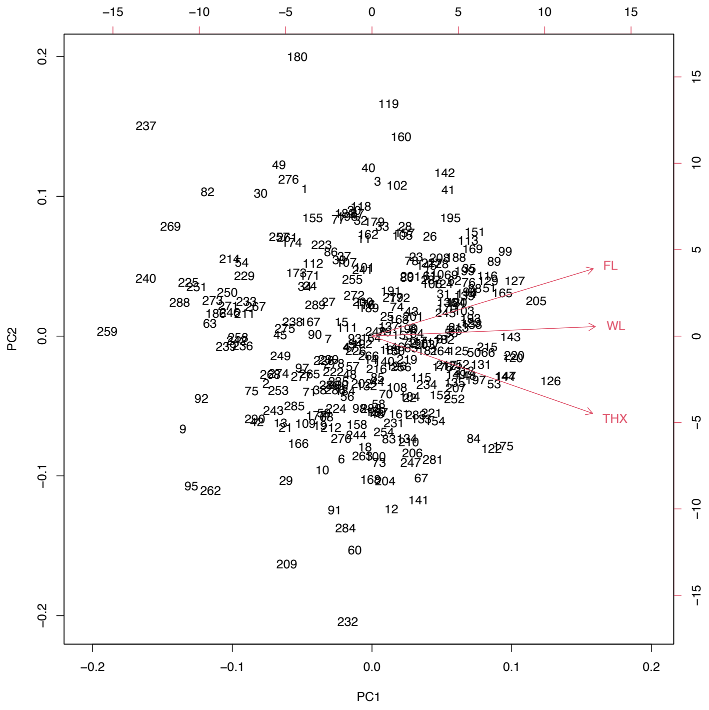


**Figure S10:** Biplot of PCA representing PC1 and PC2, as well as the contributions of each measured trait

**Statistical analyses.** All statistical analyses were conducted in R version 4.3. (2). As only approx. 50% of offspring were used in the parentage analysis, the estimate for the actual number of offspring was extrapolated from the number of assigned offspring by the parentage analysis and the overall proportion of offspring assigned for that specific plate, and then rounded to the nearest integer. Visualizations were created using *ggplot2* (3) and/or *interactions* (4). Stratified bootstrapping was done using the package *sampling* (5). If not stated otherwise, linear and generalized linear mixed models were conducted using the package *lme4* (6) with experimental units as a random effect. In analyses with copulations as the level of observation, the male and female IDs were also included as random effects.

Negative binomial models (count data) or observation-level random effects (binomial) were used where necessary to control for overdispersion (7). Arcsine-square root transformation, log transformation, and scaling and centering were used on variables where necessary to meet model assumptions. We removed interactions from models where $P > 0.1$.

Bateman gradients, i.e. the slopes between individuals' numbers of mates and the resulting offspring, were generated by fitting a generalized mixed-effects model [negative binomial *glmmTMB* (8)] controlling for overdispersion, using the replicate ID as a random effect and the number of mates, sex, OSR and density with all interactions as fixed effects. Only individuals with at least one offspring were included in the analysis to enable comparisons of slopes between the sexes (9).

The opportunity for sexual selection ($I_s$), or the variance in relative fitness, is a standardized metric of the difference in the potential for sexual selection on either sex (10). $I_s$, the ratio between the variance and the squared mean of the number of offspring, was calculated for each treatment combination and sex separately, pooling individuals across the five replicates. Confidence intervals, based on percentiles, were bootstrapped (n = 1000).

To obtain linear selection gradients with comparable results, male traits of interest (thorax, wing, and foreleg length) were standardized to ~$N(0,1)$ within each experimental unit. Additionally, we used PC1 as male composite body size. Within each unit, the relative fitness of males was calculated by dividing each male's total offspring by the mean number of offspring across all males in the unit. Selection gradients were calculated for all measured phenotypic male traits by fitting linear regressions for level of OSR and density, with relative fitness as a response variable and the standardized trait as a predictor. Statistical significance of differences between slopes was determined by the two-tailed distribution of *z*-scores ($z = \frac{slope_1 - slope_2}{\sqrt{{SE_1}^2 + {SE_2}^2}}$).

The relative contributions of mating success (MS), fertilization success (FS), and fecundity (Fec) to the overall reproductive success of males were calculated by variance partitioning. As estimates of FS and Fec were limited to instances of MS > 0, missing values were addressed by supplementing FS and Fec with their respective mean values for each unit. This approach prevents underestimating the contribution of MS while maintaining the inherent variance structure (11). Variance associated with each population factor (OSR, density) was extracted from the differences in the marginal $R^2$ values, obtained from stepwise regression. Confidence intervals for the extracted variances were bootstrapped (n = 1000), stratified by experimental unit, with a stepwise regression conducted for each resample.

For our analysis of fertilization success, we used an adjusted value of post-copulatory success (PCS), depending on the number of sperm competitors (12): $adjusted\ PCS = \ observed\ PCS * \frac{N\ competitors - 1}{observed\ PCS*(N\ competitors - 2)} + 1$. For males with >1 matings, the mean of the adjusted PCS across all their matings was used. We also used the term “remating interval” to describe the average time between a male’s mating with a female and her next mating, i.e. how long a male was able to delay a mating on average. However, as this was quantifiable only for those matings that preceded another one within the time frame of this experiment, we also used the “overall last-male”, i.e. proportion of females for which the male is the last male until the end of the experiment (11). Notably, the remating interval could not be calculated for males that were in the last-male position in all their matings. Thus, the longest remating intervals were not captured in this variable.

Finally, we used piecewise structural equation modelling to conduct our structural equation models using the package *piecewiseSEM* (13). For the model on overall reproductive success (“Fitness”) with mating success and fertilization success as predictors, we used a version of the fertilization success variable that included zeroes for those individuals that did not mate to avoid excluding those individuals from the analysis and so underestimating the effect of mating success. The models were visualized using the “neato” layout in the package *DiagrammeR* (14).
